# Unbiased learning of protein conformational representation via unsupervised random forest

**DOI:** 10.1101/2024.11.30.626148

**Authors:** Mohammad Sahil, Navjeet Ahalawat, Jagannath Mondal

**Affiliations:** Tata Institute of Fundamental Research Hyderabad, 36/P Gopanapalli Village, TS-500046, India; Department of Bioinformatics and Computational Biology, College of Biotechnology, CCS Haryana Agricultural University, Hisar 125 004 Haryana, India

## Abstract

Accurate data representation is paramount in biophysics to capture the functionally relevant motions of biomolecules. Traditional feature selection methods, while effective, often rely on labeled data based on prior knowledge and user-supervision, limiting their applicability to novel systems. Here, we present *unsupervised random forest* (URF), a self-supervised adaptation of traditional random forests that identifies functionally critical features of biomolecules without requiring prior labels. By devising a memory-efficient implementation, we first demonstrate URF’s capability to learn important sets of inter-residue features of a protein and subsequently to resolve its complex conformational landscape, performing at par or surpassing its traditional supervised counterpart and 15 other leading baseline methods. Crucially, URF is supplemented by an internal metric, the *learning coefficient*, which automates the process of hyper-parameter optimization, making the method robust and user-friendly. URF’s remarkable ability to learn important protein features in an unbiased fashion was validated against 10 independent protein systems including both both folded and intrinsically disordered states. In particular, benchmarking investigations showed that the protein representations identified by URF are functionally meaningful in comparison to current state-of-the-art deep learning methods. As an application, we show that URF can be seamlessly integrated with downstream analyses pipeline such as Markov state models to attain better resolved outputs. The investigation presented here establishes URF as a leading tool for unsupervised representation learning in protein biophysics.

---

Accurate data representation is critical for extracting meaningful insights from data, especially in protein biophysics via molecular dynamics (MD) simulations.^1,2^ These simulations, which offer detailed atomistic views of biological processes, involve thousands of degrees of freedom, not all of which are relevant to key biochemical properties like drug binding affinity.^3^ Irrelevant degrees of freedom can introduce noise, complicating the analysis.^4^ Thus, there is a pressing need for robust methods that can selectively capture the most functionally relevant protein motions. ^5,6^

Recent advances in machine learning, particularly unsupervised learning, have spurred significant progress in representation learning from high-dimensional simulation data. Two primary approaches: dimensionality reduction and feature selection, have been employed to this end. Initial efforts focused on linear dimensionality reduction techniques, such as principal component analysis (PCA)^7^ and time-lagged independent component analysis (TICA),^8^ which assume that larger variance or slower motions are indicative of functional relevance. Introduction of non-linear methods like autoencoders^9^ and its further refinement with variational^10^ and time-lagged^11^ modifications have introduced greater flexibility, and increased ability to learn non-linear conformational changes. But their reliance on reconstruction error as a measure of representation quality and requirement of fine-tuning of hyper-parameters has proven problematic. In fact, reconstruction error has been found to be unrelated to representation quality,^12^ hence necessitating atleast partial supervision.^10,13–15^ Reconstruction-free deep learning approaches, such as deepTICA^16^ and VampNets,^17^ have emerged as alternatives, optimizing time-lagged eigenvalues or VAMP2 scores instead. While these methods offer improvements, they are still grounded in dimensionality reduction, which assumes that reducing to a lower-dimensional latent space will capture all relevant protein motions. This assumption is not always valid, as all high-dimensional inputs including noise contribute to the low dimensional latent representation.

In contrast, feature selection approaches aim to identify specific degrees of freedom that are directly responsible for functional conformational changes, while discarding others that might contribute to noise or irrelevant variation.^4,18^ This selective process can be implemented through various strategies. Filter methods, for instance, evaluate individual features based on their correlation with biochemical properties or statistical measures, such as mutual information or VAMP2 scores, to isolate features most indicative of functional dynamics.^19^ Wrapper methods, such as AMINO,^20^ MoSAIC^21^ and spectral oasis, ^22^ go a step further by selecting combinations of features that collectively explain protein function, often by using iterative testing or optimization to identify the most informative subset. Meanwhile, embedded approaches leverage algorithms like random forests^23,24^ or neural network-based saliency maps^25^ and integrate feature selection directly within the training process. These embedded methods extract features by analyzing feature importance scores generated from models trained on biochemical properties, focusing on the degrees of freedom that the model deems essential to its predictions. However, most of the methods are supervised or semi-supervised methods, which require prior knowledge about the system of interest. Such methods, while effective, limit the exploration of novel systems where labels are unavailable. The challenge, therefore, is to develop an unsupervised method that can identify functionally relevant features without relying on predefined labels.

In this work, we address this knowledge gap by introducing unsupervised random forests (URF) for feature selection in protein biophysics. Unlike traditional random forests, which are supervised, URF can learn the internal structure of the data and identify the most informative features without the need for labels. This makes URF particularly suited for uncovering the critical degrees of freedom that govern protein function, enabling more accurate and noise-free data representation. Although URF was originally proposed alongside supervised random forests for limited tasks like clustering or outlier detection, ^26,27^ it has not gained widespread traction potentially due to extremely large memory requirement. Here, via introducing a more memory-efficient implementation, URF is repurposed for feature selection, marking its first application in protein biophysics or representation learning. Through extensive benchmarking against standard protocols, supervised random forests, and existing state-of-the-art baseline approaches, URF demonstrates performance better than existing approaches, highlighting its potential as a powerful tool for unsupervised representation learning. Importantly, URF learned representation is functionally meaningful and facilitate downstream analysis like building Markov models.

## Results

### Unsupervised selection of protein’s representation via URF

From molecular dynamics (MD) simulations, a large set of internal coordinates-such as native contacts, or torsional angles were estimated to define process(s) of interest, such as conformational dynamics, bio-molecular recognition, or both. The MD data, *X*^*MD*^ (*t_trajs_, t_steps_, f*), consists of independent simulation replicates (*t_trajs_*) with *f* -dimensional features or degrees of freedom (descriptors). A concatenated *X*^*MD*^ was used as the input features *X* (*n, f*), where *n* is the total number of data instances 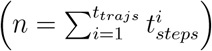.Unlike traditional supervised random forests that rely on predefined labels for classification, URF approach, as presented here, does not require any apriori labels. Instead, synthetic data (*X*^*′*^) was generated from *X* as (Fig. 1):

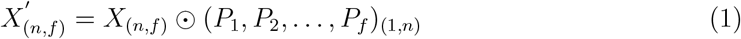

where *Pi* are independent permutation matrices that randomly shuffle different features of the input *X* independently. These random permutations disrupt the internal structure of *X* (e.g., covariance between features), ensuring that *X*^*′*^ differs from *X* in its internal structure. Ideally, *X*^*′*^ is of the same size as *X* and is referred to as permuted data. Alternatively, *X*^*′*^ can either be *marginal*, where data instances from *X* were selected randomly while maintaining similar margins hence closely resembling *X*; or *random* where data instances were randomly chosen from *X* with replacement.^28^ A *fictitious* synthetic dataset, which had no relation to *X* but was structured similarly, was also generated as a negative control.

**Figure 1:**
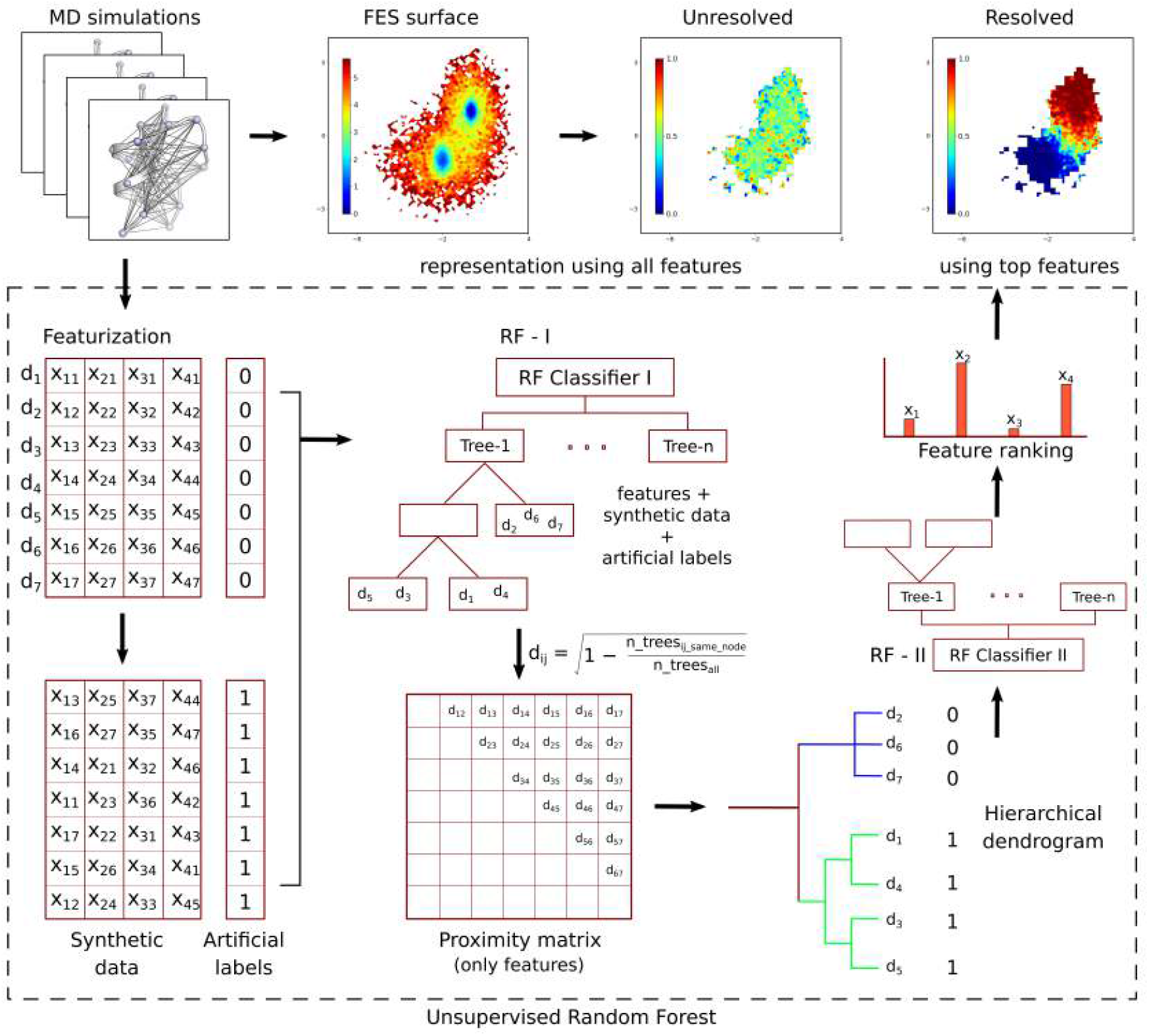
URF: The schematic of unsupervised random forest pipeline for resolving the protein’s conformational representation.

A combined dataset of *X* + *X*^*′*^ is then labeled as 0 or 1, where these labels are artificial and intended to trick the random forest algorithm into performing its standard supervised classification approach:

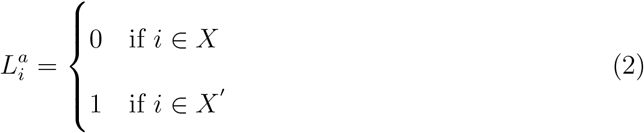

A pseudo-random forest (RF-I) is trained on the combined data (*X* + *X*^*′*^) and *L*^*a*^ to learn the internal structure of *X* by contrasting it with *X*^*′*^ . In a typical random forest estimator (*T*), the data subset is progressively split into deeper nodes until leaf nodes contain only pure samples from *X* or *X*^*′*^ . By tracking arbitrary pairs of data points from *X* in the successfully trained RF-I, one can determine whether these pairs lie within the same or different leaf nodes (pictorially represented in Fig. 1). The distance between two such data instances is then defined as:

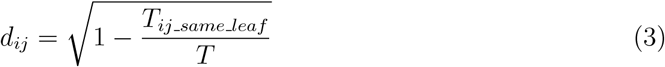

where *Tij same leaf* is the number of estimators where data instances *i* and *j* are in the same leaf node, and *T* is the total number of estimators. The square root reduces the impact of outliers.

In traditional random forests used for supervised classification, a limited number of estimators (typically tens to hundreds) is sufficient, and tree depth is restricted to avoid overfitting. However, for URF, RF-I should consist of a much larger number of estimators *T* with no restrictions on tree depth, as over-fitting is not a concern. While ideally, *n* − 1 estimators would be required to fully resolve all data pairs, a five-fold cross-validated RF-I with 1000 trees performs satisfactorily.

Using the *dij* derived from RF-I, an agglomerative hierarchical dendrogram is built for *X*, with branch distances calculated as the average of all data pairs. A new set of hierarchical labels (*L*^*hc*^) is then assigned by dividing the final dendrogram into the desired number (*nhc*) of branches. Finally, a second random forest (RF-II) is trained on *X* and *L*^*hc*^.

During RF-II training, a non-leaf parent node with *nparent* data instances is split into two child nodes based on the feature that provides the highest information gain (Δ*G*) from an arbitrary estimator with *ntree* data instances. To minimize unnecessary splits, the information gain is calculated for each available feature, and the weighted information gain is assigned as an importance score to the feature selected for the split.

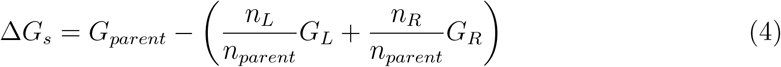

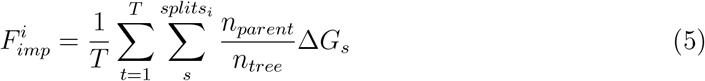

where *GL* and *GR* are the child nodes with *nL* and *nR* data instances, respectively, and 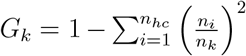 is the Gini impurity. The feature importance scores are normalized so that their total sum equals 1. Ultimately, the output of URF is a set of *f* feature importance scores, which are used to select the top features that best describe the input *X*.

### Error estimation

For RF-I and RF-II, the input data is five-fold cross validated into 0.7:0.3 train-test sub-samples. For RF-I, equation 3 (T estimators) iterates over all cross validations (*ncv*). For RF-II, five trained models were treated separately and deviation in results were used as a measure of error.

### Learning coefficient

A careful assessment indicate that efficiency of URF can be detected at hierarchical dendrogram and RF-II steps, even though sub-optimal hyper-parameters were used at other steps. For *L*^*hc*^ composed of *n_hc_* classes, let 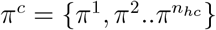 be a set of its populations. A cutoff (*n_cut_*) for outlier classes can be defined to identify outlier (*n*^*b*^) and non-outlier classes (*n*^*a*^):

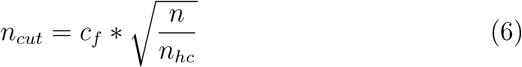

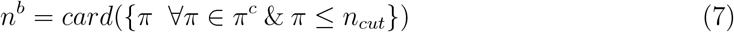

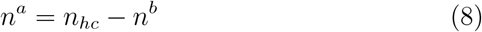

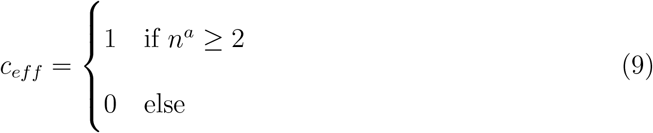

where a manually curated cutoff factor (*c_f_* = 2) was used to define *n_cut_*, and *c_eff_* is binary metric for goodness of hierarchical dendrogram.

For RF-II trained on *X*^*train*^ + *L*^*train*^ subsample of *X* + *L*^*hc*^, a *n_hc_* × *n_hc_* dimensional confusion matrix (*C*) was generated on unseen subsample (*X*^*test*^ + *L*^*test*^). Utilizing definition of *f*1 score 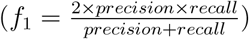,*n_hc_* accuracy values were measured for every class of *L*^*hc*^:

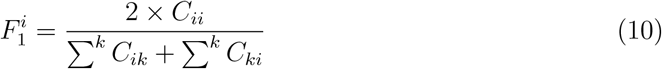

A *learning coefficient* (*LC*) was estimated by combining efficiencies of hierarchical dendrogram and RF-II:

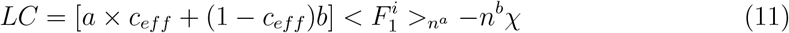

where a and b are upper and lower (subjected to *χ*) bounds for *LC* and *χ* is penalty for outlier classes. For a=1, b=0 and *χ*=0, *LC* can be:

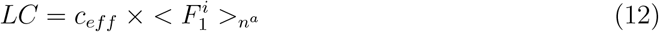

In principle LC can have 0-1 values, but practically observed only either close to zero or close to one which corresponds to unsuccessful/successful training of URF.

### Memory efficient implementation

Though individual steps of random forest and hierarchical dendrogram are already implemented in scikit-learn,^29^ and fastcluster, ^30^ a direct-user-friendly implementation is lacking potentially due to its exponential memory requirements. For *X* of *n* samples, equation 3 outputs an array of ^*n*^*C*_2_ elements. For n=154k, around 11.8 billion elements needs to be saved which even in condensed form requires ∼88 GB and a total of ∼120 GB of working memory. The MD data (*X*) is generally of order 10^6^ or higher and can require upto tera-bytes of working memory. Using part of data can lead to loss of efficiency. Therefore, three different implementation schemes were designed with extensive benchmarking to avoid loss of efficiency and default hyper-parameters estimation (Extended Data Fig. 1), (a) *efficient* : a uniform sub-fraction of *X* (at least 3*n_cv_T* data instances) is used in equation 3 and subsequent steps. Depending on sub-fraction used in equation 3, this schemes significantly reduces both time and memory requirement and maintains the URF efficiency. (b) *fit-predict* : the input *X* is used in two parts; a smaller fit part (at least 3*n_cv_T* data instances) on which equation 3 is estimated via efficient algorithm up to *L*^*hc*^, and a predict part (remaining data) for which *L*^*hc*^ is determined via estimating equation 3 in batches and a knn-like algorithm with *n_cut_* neighbours. This scheme has significantly reduced (larger compared to efficient) time and memory requirement but uses complete *X* and maintains URF efficiency, and (c) *low-mem*: equation 3 is estimated via on-the-fly batches of *X* without saving complete output. This scheme utilizing modified nearest-neighbour-chain algorithm^31^ uses complete *X* with minimal memory requirement but is extremely slow.

### URF resolves proteins’ conformational landscape

The efficacy of the URF in distinguishing the ligand-bound conformation from the unbound conformation was evaluated on a previously reported MD simulation trajectory (supplementary methods, data-1).^32^ The T4 lysozyme (T4L)-benzene, a well-characterized proteinligand system, exhibits distinct ligand-dependent conformations for both bound and unbound states. This simulation involved unbiased binding process where benzene starts in an unbound state and spontaneously locates its binding pocket within T4L. During the transition from the unbound to bound state, T4L is expected to adopt state specific conformations. To detect these conformations specific to the bound and unbound states, a set of features was defined based on the native contacts within T4L. By enumerating the minimum distances between residue pairs with *Cα* distances ≤ 10°*A*, 755 native contacts were used to define the protein’s conformation at any given instant. Initially, we generated a TICA-derived free energy surface (FES) of the protein conformations, annotated by the probability of observing a bound state on the FES. This was to determine whether the bound state occupies a discrete and distinct location on FES (methods).^23^ As shown in Fig. 2A, the bound state is intermingled with the unbound state (no classifier), suggesting either the absence of specific conformations for the bound state or a limitation in the analytical pipeline.

**Figure 2:**
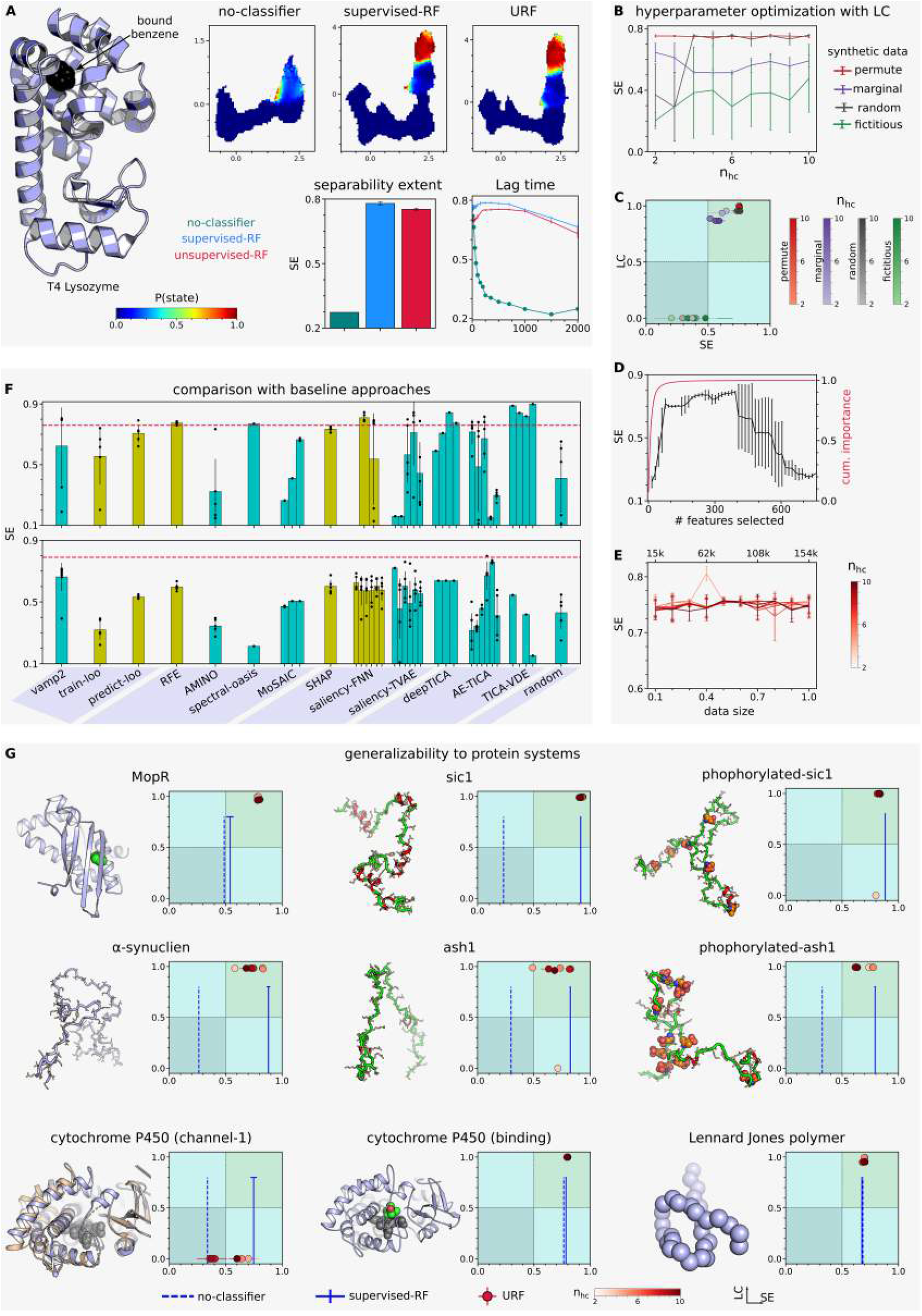
URF learns conformational representation: (A) Representation plots of native contacts for bound state of T4L via different schemes, quantified by separability extent (SE) and different lag times. (B, C) URF performance for different *X*^*′*^ and *nhc* settings and its correlation with LC. (D) Varying URF performance and cumulative importance score with increasing number of selected features. (E) URF performance with decreasing data size (fraction of total data, number of data points) with ‘efficient’ scheme. (F) Comparison of URF with baseline approaches for T4L (top) and mopR (bottom). Dashed line indicate URF, color of bar indicate supervised (olive) or unsupervised (darkcyan) nature of approach. (G) Applicability of URF for different systems. (n=5, except vamp2 approach in (F) where n=6 (T4L) and n=10 (mopR). Most error bars in (G) are small and not visible.)

To refine the analysis, 755-dimensional input data was binarily labelled based on prior knowledge of bound and unbound states. Upon successful training, the supervised random forest (RF) model identified 200 native contacts that collectively contributed to a cumulative importance score of 0.801±0.001. When the FES was generated using the selected native contacts and annotated with the probability of finding a bound state, the resulting representation displayed a distinct, exclusive region corresponding to the bound state. However, this outcome was expected, as the supervised RF model was trained using prior knowledge of the bound and unbound states, a condition that would not apply in an ideal scenario.

To address this limitation, unsupervised random forest (URF) protocol was applied which does not require any prior information. Using the URF-selected top 200 native contacts, we again observed a well-resolved FES for the bound state (Fig. 2A). The resolution was comparable with the supervised RF approach as measured by separability extent (SE, methods). In a two-state system like T4L, resolving the bound state automatically implies a similar resolution of the unbound state (Supplementary Fig. 1). This observation was consistently reproducible across other independent trajectories, as reported in the previous work. ^23,32^ Although the TICA-based FES is fundamentally dependent on the chosen lag time, ^33^ the URF-based resolution of the protein’s conformational landscape demonstrated robustness across the range of lag times tested (Fig. 2A, Supplementary Fig. 1)

#### Guiding hyper-parameters optimization using Learning coefficient(LC)

Similar to any machine learning algorithm, the performance of URF is sensitive to suboptimal hyper-parameter choices in its multi-step protocol. Out of rigorous search, URF efficiency significantly depends on varying synthetic data type (*X*^*′*^, permute, marginal, random, fictitious) and hierarchical labels (*n_hc_*, 2-10) (Fig. 2B). Here learning coefficient (LC) can internally evaluate the effectiveness of the URF protocol, with the best-performing hyper-parameters identified based on the highest LC values (Fig. 2C). For instance, the high and low performance of URF with permute and fictitious *X*^*′*^ across all *n_hc_* corresponds to high and low LC values respectively. The LC was sensitive enough to detect small changes in performance (marginal with varying *n_hc_*) or hyper-parameters (random with *n_hc_* = 3 − 4), hence can guide URF optimization without apriori knowledge.

Beyond the URF protocol, the results were also sensitive to the number of top features selected for downstream analysis. In this context, the cumulative sum of importance scores guided hyper-parameter selection. As the number of features increased, the SE initially rose and then plateaued as the importance score saturated (Fig. 2D, Supplementary Fig. 2). Beyond this saturation point, SE began to decline as more unimportant features were added without a corresponding increase in the importance score. Therefore, the optimal number of features should be chosen near the saturation point in the importance score, which is also guided by LC (Extended Data Fig. 2).

The URF protocol’s high memory requirement is driven by the number of data instances involved in proximity estimation (equation 3). However, this can be significantly reduced using an ‘efficient’ scheme without compromising performance (Fig. 2E, Supplementary Fig. 2). It is emphasized that the URF protocol defined here is not equivalent to supervised learning on labels derived from straightforward clustering. Unlike URF, direct hierarchical clustering, k-means (euclidean), or k-medoids (euclidean, proximity) are not stable with varying *n_hc_* (Extended Data Fig. 3), and the LC applied to these algorithms often yields misleading results.

**Figure 3:**
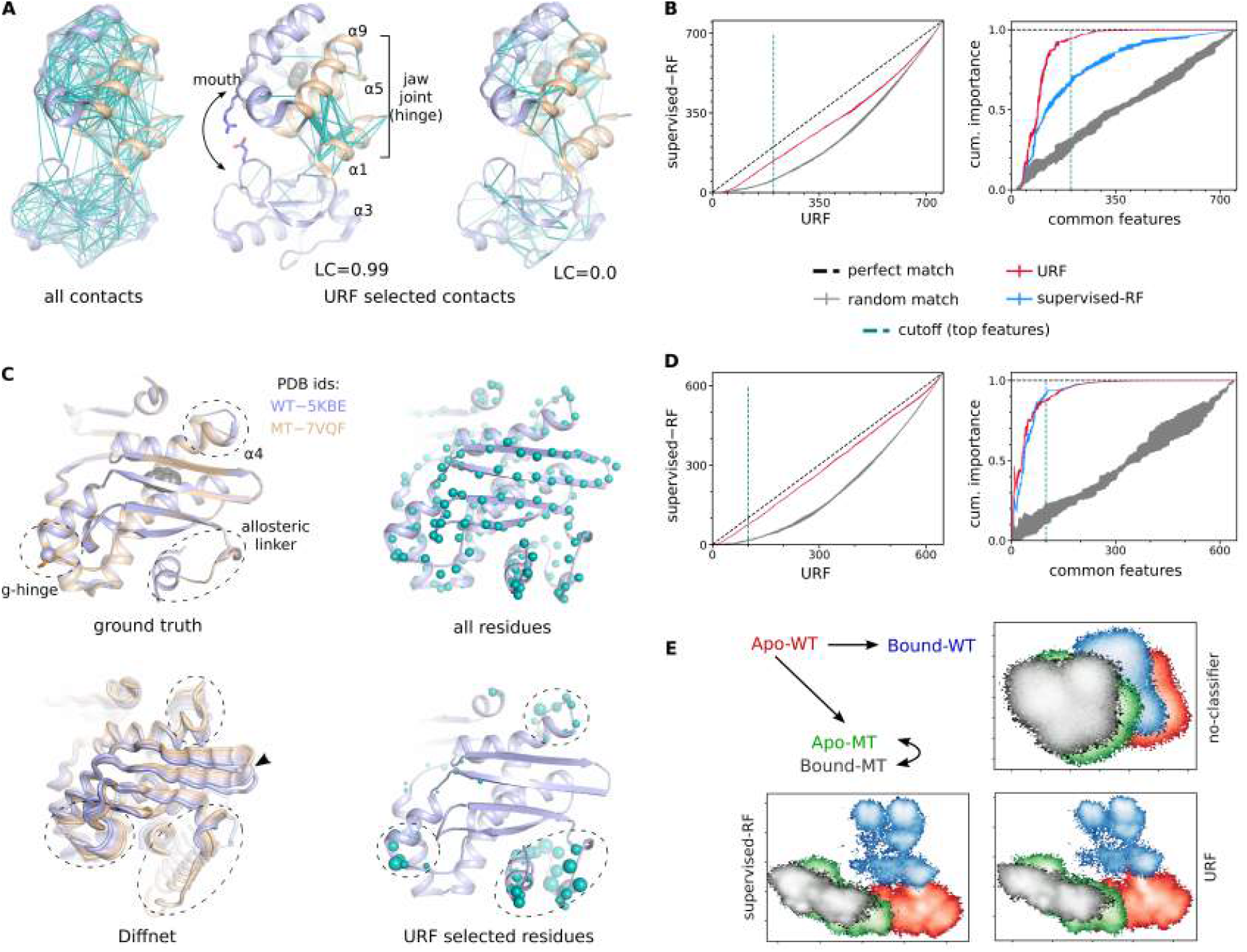
URF identifies functional regions: (A) Pictorial visualization of native contacts as lines joining residue pairs. All and URF selected native contacts were shown in T4L. Width of line corresponds to URF derived importance scores. LC values indicate learned URF efficiency. The residues in sticks are E22 and R137. (B) Quantitative similarity of URF selected features with supervised-RF for T4L. (C) Important functional regions in mopR determined via crystallography (ground truth), URF and DiffNet. The g-hinge region involving dynamic allostery was not observed in crystals, but identified via simulations and mutagenesis ^41^ (Supplementary Fig. S7A). The dashed circles and arrowhead indicate successfully identified and false positive functional regions respectively. (D) Quantitative similarity of URF selected features with supervised-RF for mopR. (E) Expected pattern and estimated FES plots of different states of mopR with no-classifier, supervised-RF and URF schemes. The error bars in (B) and (D) merged together and are visible as width of the curves (n=5).

#### Assessing URF across diverse systems

The applicability of URF protocol was explored on diverse systems encompassing folded to disordered proteins to beads-on-a-string polymers (Fig. 2G). All these systems were previously characterized for their functional states (ground truth) via both simulations and experiments by our and other research groups. For instance; a candidate-in-design phenol biosensor mopR^34^ exhibits a crucial intermediate state for ligand selection, ^35^ cytochrome P450 has state-specific conformations depending on ligand binding or channel-1, ^36^ intrinsically disordered protein Sic1, Ash1, *α*-synuclein exhibit compact and expanded conformations here demarcated by their experimental Rg values, ^37,38^ and LJ polymer also exhibit conformations ranging from compact to expanded states as per simulation observed Rg values ^39^ (detailed in supplementary methods).

The efficacy of URF to resolve these functionally crucial states was probed, specifically the statistically minor functional i.e. rare states . URF was tested for *n_hc_*=2-10 to encompass its hyper-parameter sensitivity (if any). Its performance was compared with no-classifier and supervised-RF serving as negative and positive controls respectively. SE was used as an explicit measure to estimate URF performance and its comparison with no-classifier and supervised-RF (x-axis in Fig. 2G plots). The URF was further evaluated for LC to determine whether this internal metric accurately reflects its performance (y-axis in Fig. 2G plots).

We observed that (Fig. 2G, Supplementary Fig. 3): a) the URF effectively resolves the representation and, in some cases (mopR), even outperforms the supervised-RF, b) for most systems and across the majority of hyper-parameters, URF outperformed the no-classifier approach and performed comparably to the supervised-RF, c) in specific instances, such as with certain hyper-parameters in systems like *α*-Synuclein and ash1, URF significantly outperformed the no-classifier approach (resolving the representation with a SE≥ 0.5) but did not match the performance of supervised-RF, d) the LC internally detects the successful learning of URF in all cases, except for two instances where false negatives were observed in two systems (ash1 and phosphorylated-sic1) and e) URF was ineffective for cytochrome P450 (channel-1), as predicted by LC=0 for all hyper-parameters and SE similar to the no-classifier approach across most hyper-parameters. In some cases for cytochrome P450 (channel-1), URF approached the performance of supervised-RF but with substantial errors, i.e., unstable performance with LC=0. Additionally, the systems cytochrome P450 (ligand binding) and a simplistic 32-bead LJ polymer achieved resolved representations even with the no-classifier approach, indicating that these datasets do not require further improvement. However, it should be noted that in an ideal scenario, this information would not be known a priori. In these cases, URF maintained the existing resolution, similar to supervised-RF and as predicted by LC. Therefore, even if the representation is already resolved and refinement is not necessary, the URF protocol does not degrade the resolution. Overall, URF can learn a resolved conformational representation with LC predicting its performance without any prior information of ground truth. Out of 198 cases tested for all systems, LC correctly predicted the URF performance for 97% cases, with only 6 cases of false negative were observed.

#### Comparison with baseline approaches

Performance of URF was compared with 13 existing approaches across both supervised and unsupervised schemes (Fig. 2F). These methods can be broadly categorized into five categories: (i) *filter* approaches (VAMP2, train-loo, predict-loo), ^19,29^ *wrapper* methods (RFE, AMINO, spectral-oasis, MoSAIC), ^20–22,29^ (iii) *embedded* approaches (SHAP, saliency-FNN, saliency-TVAE), ^25,40^ (iv) *deep learning improvements* of TICA-based representation (deepTICA, AE-TICA, TICA-VDE),^10,16^ and (v) a negative control involving random feature selection. Although the deep learning methods are not feature selection protocols per se, they have been utilized in resolving the TICA based FES. For some approaches, such as saliency-TVAE and deepTICA, multiple hyper-parameter choices were possible, hence their results are represented by multiple bars in Fig. 2F. Cross-validation error was not applicable for methods like TICA-VDE and deepTICA performs its internal time-dependent split. Additionally, AMINO estimates the number of features to be selected, and MoSAIC provides clusters of selected features, while other methods do not specify this, so number of selected features were maintain same as URF. Specific details on these approaches are provided in the supplementary methods and Supplementary Figs. 4 and 5.

**Figure 4:**
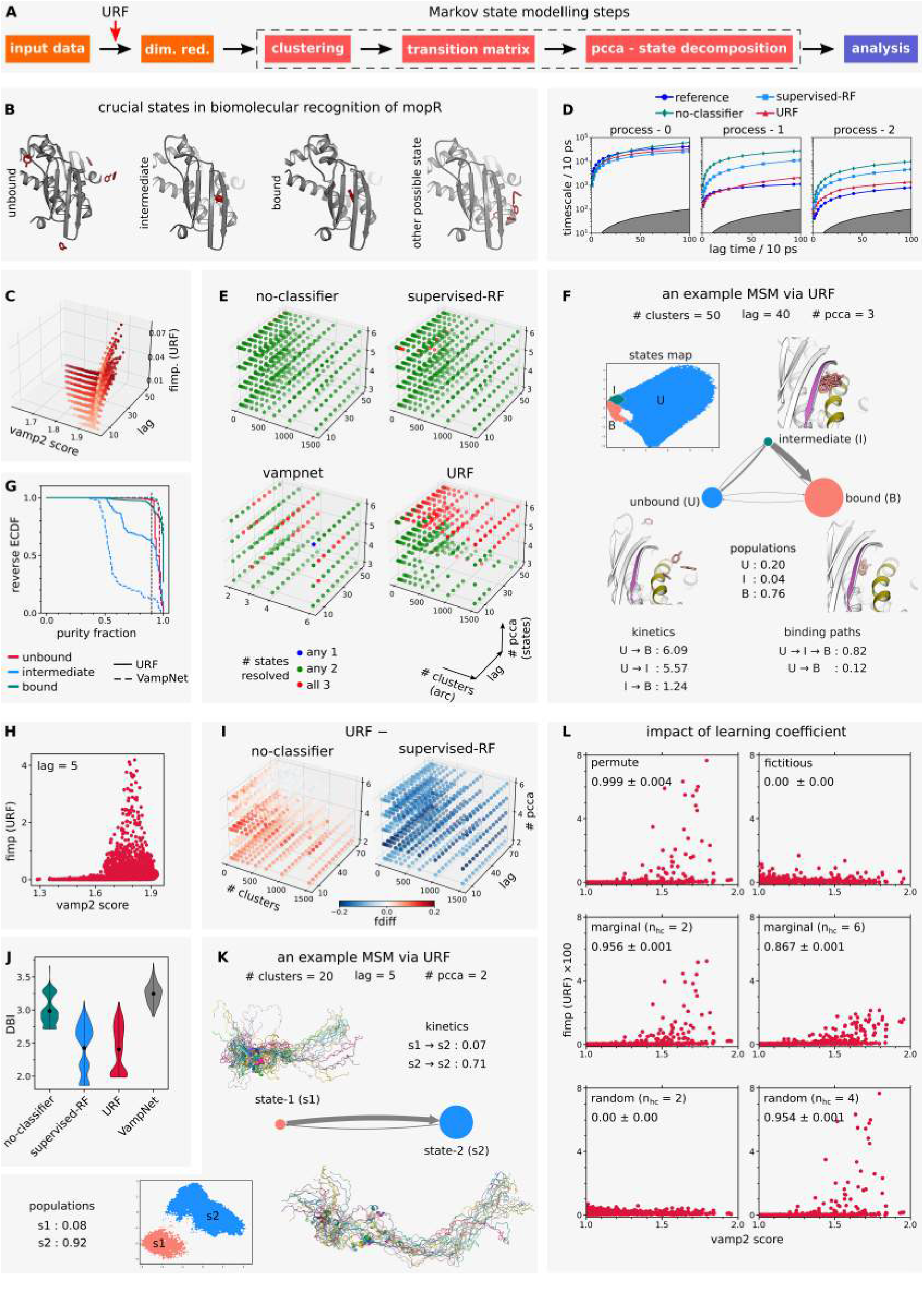
URF facilitate downstream analysis: (A) Proposed integration of URF at the start in MSM pipeline. (B) The three crucial binding states of mopR, i.e., unbound, intermediate and bound states. An example of other possible state is also shown. These representative frames were drawn based on prior knowledge. ^35^ (C) Relation between VAMP2 and URF derived fimp at varying lag times for mopR. (D) Comparision of top three implied timescales derived from different schemes with reference handcrafted MSM. ^35^ (E) Number of crucial states resolved in markov models build via different schemes for mopR. Each dot represent a markov model and color represent the number of resolved states. *arc* means architecture of VampNet (methods). (F) An example 3 state MSM built via URF scheme for mopR. (G) The purity of three states of successful 3-state markov models in URF and VampNet (red dots in E subplots). (H) Relation between VAMP2 and URF derived fimp for *α*-synuclein. (I,J) Comparison of markov models built via different schemes for *α*-synuclein. Larger *fdiff* and smaller *DBI* indicate better resolved models. The values represents mean (n=5). (K) An example MSM derived via URF scheme for *α*-synuclein. The representative snapshots are most expanded frames of each metastable state. (L) The VAMP2-fimp (URF) relation for T4L at different hyper-parameters with varying LC values. (The values represent mean for VAMP2 scores, errorbars not shown for clarity.)

The benchmarking was conducted on two protein-ligand binding systems: T4L/benzene and MopR/phenol. These systems had extensively long in-house MD simulation trajectories that capture ligand-binding events in real time, making them suitable for comparing the ability of each methods to resolve ligand-bound versus ligand-unbound protein conformations. ^32,35^ For all approaches, the representation on selected features or learned latent representations was assessed via SE. Fig. 2F provides a quantitative comparison of these approaches with URF in terms of SE. The assessment yielded several key observations: (a) URF is either superior to or on par with the base-line approaches; (b) it is unsupervised than approaches like SHAP, saliency-FNN, and predict-loo, which perform comparably to URF; (c) it is more generalized compared to methods like TICA-VDE, which outperforms URF for T4L but performs poorly for MopR; (d) it is self-guided by the LC, unlike methods such as saliency-TVAE, where the learning curves do not correlate with performance, and most other approaches (e.g., SHAP, AMINO, spectral-oasis) that lack a metric of goodness; and (e) it exhibits significantly lower error bars compared to methods like saliency-TVAE.

### URF extracts functionally meaningful representation

For proteins, especially folded, not all residues contribute to function or undergo conformational changes. Instead, a small but critical subset of residues can explain these functions, and various methods such as DiffNet have been developed to identify them. ^13^ Given that URF-selected representation is resolved for functional states, it is reasonable to guesstimate that URF might be learning the functionally relevant regions of a protein. In this regard, URF was probed whether its learned features corresponds to functionally important residues, i.e., structural features estimated for all residues were used as input to URF and probed if functionally important residues are attributed with higher importance scores. This analysis was performed for T4L ^32^ and mopR ^41^ in light of their known ground truths and via comparison with supervised-RF and baseline approaches.

T4L broadly constitutes N (tail) and C (ligand binding pocket) terminal domains, making an arc shaped structure connected via a central hinge (jaw-joint) and forming a mouth between two domains (Fig. 3A). As a function of ligand binding, conformational motions at the hinge region lead to opening/closing of mouth region, which can be detected via salt bridge change between E22-R137 or D20-R145 residue pairs.^42,43^ We probed whether URF is able to learn this functionally relevant hinge region. All the 755 native contacts i.e., features (data-1) spanning the complete structure of T4L were used as input to URF. The selected native contacts were visualized by reweighing with the importance scores learned by successfully trained URF (LC=0.99). The native contacts assigned with highest importance scores were localized in the functionally important hinge region (*α*1, *α*5 and *α*9), i.e., URF has learned functionally relevant residues in T4L (Fig. 3A). It is emphasized that all native contacts with high importance scores are localized in a discrete region (hinge), an indication that URF is able to learn targeted conformational changes in protein. As a contrast, an unsuccessfully trained URF (LC=0) neither attributes significant importance scores to hinge nor it localized to any other discrete region. This observation also emphasizes that LC can internally detect the learning of functional regions.

As a comparison with supervised-RF, URF learned similar native contacts but with differences in absolute values of importance scores (Fig. 3B). For instance, out of top 200, URF selects 141±2 native contacts common with supervised-RF in contrast to 57±3 achievable by random selection. A significant portion of URF-selected contacts differed from supervised-RF, but the most important contacts located in hinge region belongs to common group and contributes most of the importance scores (URF: 0.95±0.01, supervised-RF: 0.68±0.02, random-selection: 0.29±0.04, Fig. 3B). It is to note that, in literature, ^42^ *α*3 is also included as crucial part of hinge region but did not come to be important in URF or supervised-RF, potentially because of differences in simulation-type, input data and mutations (this work utilizes L99A mutant of T4L).

As a second system, an NtrC transcriptional regulator and phenol biosensor, namely, mopR^34^ exhibits multiple functional allosteric regions, ^41,44,45^ (a) allosteric-linker: an allosteric effector region which transfers the signal of phenol detection to downstream domains, (b) g-hinge: this region allosterically controls the binding pocket and a G148P mutant in this region led to 7-fold increase in binding affinity, and (c) *α*4: this region controls the downstream transcriptional activation of phenol-catabolic genes. The previous simulation-experiment-crystallography^41^ works involved a manual and tediously supervised curation of thousands of hydrogen bonds in a network leading to identification of above functional regions and their interconnections (Fig. 3C, Supplementary Fig. 6A). Here we checked if the URF can unbiasedly identify these functional regions. The input dihedral space (*ϕ, ψ, χ*1 dihedrals of all residues) from four MD simulation ensembles (data3-6, apo-WT, bound-WT, apo-MT, bound-MT (apo and bound means with/out phenol, WT and MT means wildtype and G148P mutant), methods) was used as the features for successfully training a URF (LC=0.99). The residues having dihedrals with high learned importance scores, if visualized on structure, corresponds to the functional regions of mopR (Fig. 3C), indicating that URF can learn the functional regions in proteins. For mopR also, the successful training of URF can be detected by LC values, as unsuccessfully trained URF models (low LC) were unable to identify functional regions (Supplementary Fig. 6B).

Compared to supervised-RF, URF identifies 79±1 out of 100 dihedrals common with supervised-RF (random selection=17±3, Fig. 3D). The important dihedrals are common in both URF and supervised-RF and contributes most of the importance score (URF: 0.88±0.01, supervised-RF: 0.92±0.01, random-selection: 0.17±0.07).

URF performance was also compared with DiffNet, ^13^ a current state-of-the-art deep learning based protocol for identifying the functionally important regions defining the functional states of protein. DiffNet takes separate simulation ensembles as input and hence compared only for mopR having four separate simulations. It learns the input features which best describes the differences in a overlapping group of simulation ensembles. The difference was probed in a pre-defined order as guided by binary hyper-parameter *act map* used here as 0,1,1,1 for apo-WT, bound-WT, apo-MT and bound-MT respectively (methods). A successfully trained DiffNet not only identified the functional regions in mopR, but also outputs the conformational motion spectrum and is better than URF in this respect (Fig 3C). However, it also identified an additional false positive region which is not known to be functionally relevant atleast yet. Another DiffNet model could identify g-hinge and allosteric linker regions but missed *α*4 region (Supplementary Fig. 6).

Finally, the meaning in representation plots of simulation ensembles lies in their relative locations which is indicative of its conformational differences. ^46,47^ For instance, the simulation ensembles possessing different conformations would have distinct location on representation plot. In case of mopR, the previous hydrogen bond network and tryptophan quenching studies^41^ found that mopR in apo-WT and bound-WT possess different conformational ensembles (effect of binding) and mutant variants (apo-MT, bound-MT, effect of G148P mutation) have yet different conformations (Fig. 3E, Supplementary Fig. 6). The PCA based representation using all dihedrals (no-classifier) were overlapped, therefore unable to indicate the conformational differences. The representation using URF selected dihedrals indicate three distinct locations for apo-WT, bound-WT and mutants (apo-WT, bound-WT) as expected, and similar to supervised-RF.

### Representation learned by URF facilitates downstream analysis

URF, with its resolved representation, should facilitate the downstream analysis, an ultimate aim of any representation learning protocol. We checked this on Markov state models^48^ (MSM), a reliable and widely used analysis protocol for MD data. The proposal is that introduction of URF step in early stage of MSM pipeline can help in refinement of input features, which in turn, would help build better markov models (Fig. 4A). This was checked for two protein systems: namely, ligand recognition process in mopR (data-2, reference MSM or ground truth is known), and conformational dynamics in *α*-synuclein (no ground truth or well defined states are known). The MSMs were built with three different schemes, i.e., no-classifier (negative control, base approach), supervised-RF (positive control) and URF (test case). URF was also compared with VampNet, ^17^ a current state-of-the-art deep learning based dimensionality reduction protocol that identifies the metastable states in the data by maximizing the VAMP2 scores. Finally, URF derived importance scores were correlated with MSM associated VAMP2 scores ^19^ for all systems, to address LC guided generalizability.

The MSM is a multi-step pipeline and sensitive to its own hyper-parameter choices at its three integral steps. While indirect ways are available for choosing hyper-parameters, but comparison based on single hyper-parameter settings can be potentially misleading. Hence, MSMs were built at large hyper-parameter space at every step of clustering, lag time, and pcca states. Overall, by testing URF and MSM hyper-parameters, a total of 16920 (mopR:7680, *α*-synuclein:9240) markov models were built to achieve comprehensive conclusions (methods). Few models, if detected for wrong hyper-parameters, were not considered. Finally, the quality of MSM were analyzed based on the realization of ultimate metastable states (methods).

As a ground truth (reference MSM) studied by combined experiments-simulations and handcrafted ^49^ MSM, ^35^ binding process in mopR undertake through three crucial stages, unbound →intermediate → bound (Fig. 4B). Any protein-ligand binding model should have unbound and ligand-bound states, while intermediate state is the most crucial state in mopR, responsible for its ligand selection process, and has also been confirmed by mutagenesis. MSMs may output additional metastable states which are not necessarily involved in process of interest or experimentally justified (fourth state in mopR and other possible states). Hence for a k macrostate (k≥3) MSM or VampNet, a correct model should have bound, unbound and intermediate states and goodness of a model shall depend on the purity of these resolved states (methods). The top three implied timescales from URF-MSM closely match with those of ground-truth reference MSM (corresponding to three states) as compared to no-classifier and supervised-RF (Fig. 4D, Supplementary Fig. 7). URF-integrated MSM models could resolve three crucial binding states at large hyper-parameter space, compared to only few models built with no-classifier or supervised-RF (Fig. 4E,F, Supplementary Fig. 8). Compared to VampNet which can also build models with three resolved states, URF is better with its 28.14% (1100/3909) successful attempts compared to 13.8 %(110/796) of VampNets. Of the successful URF-MSM models, most of the bound and unbound states were resolved with a purity of *>* 90% as was the case with VampNet also. The rare intermediate state was the delicate to resolve with *>* 90% purity. Around 60% of URF-MSM have intermediate state with *>* 90% purity, compared to 12.5% of VampNet (Fig. 4G). The successful building of better URF-MSM was also valid for cross-validation replicates and different hyper-parameter of URF (Supplementary Fig. 8). As a second case study, the intrinsically disordered protein (IDP) *α*-synuclein lacks well-defined conformational states, meaning there is no established reference MSM to serve as ground truth. Consequently, MSM quality was assessed using independent metrics designed to evaluate the exclusivity of metastable states. Here, exclusivity refers to the grouping of similar conformations within the same metastable state, while distinct conformations should reside in separate metastable states. Conversely, in an inaccurate MSM, distant conformations would be grouped within the same metastable state, leading to mixed metastable states. This exclusivity was quantified by higher *fd-iff* or lower *DBI* values (methods). For *α*-synuclein, the URF-MSM outperformed the no-classifier approach but did not match the performance of the supervised-RF (Fig. 4I-K, Supplementary Fig. 9). In models with higher state counts and longer lag times, URF-MSM did not surpass the no-classifier approach, likely due to the rapid conformational dynamics typical of IDPs. URF-MSM also significantly outperformed VampNet, which did not improve upon the no-classifier approach (Fig. 4J, Supplementary Fig. 9), suggesting limitations of VampNet based on data type or protein type. Though there is no ground truth for IDPs like *α*-synuclein, these findings suggest that building MSM on a selectively reduced feature space yield superior models.

VAMP2 score is well attested and routinely used with MSM and assigned to input data for its potential usefulness for building markov models. ^4,19,50^ With URF-MSM performing better than regular MSM (no-classifier), the URF outputted importance scores (fimp) were checked for their correlation with VAMP2 score. An intriguing yet expected pattern emerges: features with high fimp have high VAMP2 scores but not vice-versa (Fig. 4C, H, Supplementary Figs. 8, 9). For mopR, this pattern was visible at all lag times, and better URF-MSM were observed at all lags. On the other hand for faster dynamics of *α*-synuclein, this pattern disappears at larger lag times where URF-MSM does not improve on no-classifier. This fimp-VAMP2 relation indicate that URF is selecting features useful for building markov models as per VAMP2, as explicitly verified for the two protein systems. To address generalizability, the fimp-VAMP2 relation was further checked on all protein systems and corroborated with LC values. Intriguingly, this fimp-VAMP2 relation was evident for all successfully trained URF models in correlation with their LC values, evidently for all systems checked (Fig. 4L, Extended Data Fig. 4). In summary, URF selected features have the potential to build better resolved markov models. It is noted that URF does not improve on kinetics of markov models, nevertheless the kinetics is only meaningful in resolved MSM.

Overall, building markov models on unbiasedly selected URF based representation can output better models. In general, we claim that URF can be integrated to any downstream analysis pipeline on MD data to attain better resolved outputs.

## Discussion

Here we repurposed unsupervised random forest (URF), a self-supervised version of traditionally known supervised random forest, to extract functionally meaningful structural representation for protein conformational ensembles. URF learns the conformational representation via feature selection route, thereby the representation based on selected features not only is resolved for functional states, but also corresponds to functionally relevant structural motifs. The workflow of URF is sufficiently streamlined for interpretability compared to deep learning based approaches where latent space are abstract which lacks interpretability and exploitable structures at the latent space, thereby allowing the identification of functional protein residues and their high order relations like allostery. Importantly the hyper-parameter tuning is self guided by LC and is internally incorporated in final implementation codes. Though URF learns functional representation as good as supervised approaches, it is emphasized that URF is a standalone protocol for representation learning and not directly comparable to supervised random forest for classification/regression tasks. Here, we propose that instead of manual selection ^4,51,52^ the URF step can be incorporated in any high dimensional downstream analysis protocol for MD data like building markov models to attain better resolved outputs. Moreover accurate feature selection is a broader problem and is widely used in many disciplines involving high dimensional input data, ^53–56^ where URF may find its additional applications.

We attempted to benchmark URF performance with most baseline approaches for feature selection. Nevertheless, a large number of feature selection approaches has been devised and benchmarking was not performed with all of them. Specifically, approaches like select k best, select from model, MRMR, MDFS, FDA, Boruta, Shapash, OMNIXAI, InterpretML, Dalex, Eli5, Relief etc have been designed. ^29,57–60^ We reason that all these approaches are variations of VAMP2, train-loo, predict-loo, RFE, SHAP, supervised-RF and saliency-FNN algorithms which have been used in benchmarking. Also, most methods can be criticized due to their supervised nature. Here, benchmarking was specifically targeted for approaches that are directly or indirectly been utilized in MD and protein associated problems. URF was found to be as good as or even outperformed most of the baseline approaches. Specifically, URF performed comprehensively on three different tasks (learning representation, identifying functional regions, downstream task of building markov models) at par or better than current state-of-the-art approaches of each task. In general, URF can be added to any downstream analysis pipeline for MD data.

From critical perspective, URF could not learn functional representation for one of the systems tested (cytochrome P450-channel-1), though LC correctly predicted it as true negative. Although, LC can predict the URF performance but some cases (6/198) of false negative results were also observed. Further, URF can learn functional representation only if input data possessed enough conformational sampling, though it shall be applicable for all algorithms. Finally, representation learning algorithms including URF requires meaningful input data to learn from. For instance, the functional states of some protein differs in terms of sidechain hydrogen bonds but the input data only contains backbone dihedrals, then URF or any other representation learning algorithms can output false positive results. In this regard, graph based modifications are being introduced very recently which can learn directly from atomic coordinates but are yet to be tested widely. ^61,62^ In future, a similar GNN-URF can be devised.

## Methods

### Curation of Molecular dynamics Simulation data

This work utilizes 14 different datasets derived from molecular dynamics simulations corresponding to 7 different protein or protein-like systems, ranging from intrinsically disordered to large folded globular proteins. The protein systems include T4-Lysozyme ^32^ (data 1), phenol biosensing protein mopR ^35,41^ (data 2-6), cell cycle proteins sic1 (wildtype - data 7, multi-phosphorylated - data 8) and ash1 (wildtype - data 9, multi-phosphorylated - data 10), ^38,63^ Parkinson’s disease associated *α*-synuclein ^37^ (data 11), cytochrome P450^36,64^ (data 12, 13) and 32 bead LJ polymer ^39^ (data 14). The total data amount to around 250 *µ*s or 1.46 × 10^7^ data instances. All these systems has been characterized for their functional states (ground truth (*L*^*T*^)) in previously published works and briefly detailed in supplementary methods and Supplementary Fig. 10. For each system, high dimensional set of collective variables (*X*^*MD*^, like native contacts, dihedral space, whitened coordinates etc.) defining a process of interest were calculated, also detailed in supplementary methods.

### No-classifier, supervised-RF and URF schemes

For no-classifier scheme, complete data (*X*^*MD*^) from MD simulations was used as input. For supervised-RF schemes, the *X*^*MD*^ was trained to learn categorical or continuous labels based on functional states (*L*^*T*^). For URF scheme, the unsupervised training was performed on *X*^*MD*^. Subsequently the shortlisted version of *X*^*MD*^, in terms of number of features with high learned importance scores, was used as input for supervised-RF and URF schemes.

### Data representation and separability extent

The 2d projection of *X*^*MD*^ was generated by time lagged independent component analysis, ^8^ a widely used algorithm for MD data which also considers dynamics, an essential component of MD simulations. The resultant projection was represented on 100×100 binned surface, where each bin is assigned an empirical probability [0-1] of belonging to a particular functional state, estimated as fraction of data instances in a bin attributed to that state. The ultimate two-dimensional representation of *X*^*MD*^ indicate whether the functional state of interest have discrete and exclusive nature or is mixed with other possible states.

For a binned surface, let *tb* be the total number of non-empty bins which can be further classified into (a) bins pure higher (*b_ph_*): bins belonging to a functional state, (b) bins pure lower (*b_pl_*): bins not belong to a functional state, and (c) impure bins (*b_imp_*), pictorially represented in Supplementary Fig. 11:

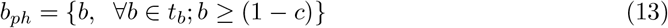

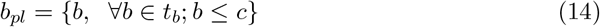

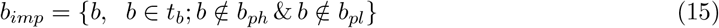

where *c* is cutoff used to define purity of bins. c=0.1 was used for all systems i.e., bins having 0.9 or above were termed as pure higher. The empirical strengths of *b_ph_, b_pl_* and *b_imp_* can be estimated as:

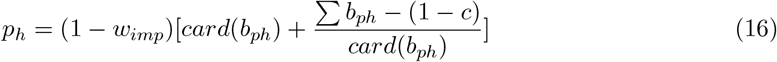

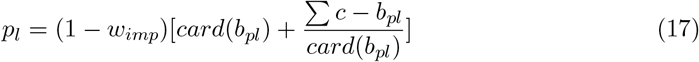

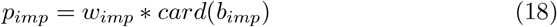

where *w_imp_* and 1 − *w_imp_* are weights associated to impure and pure bins respectively. *card* is cardinality or count function. Finally, separability extent (SE) was estimated as:

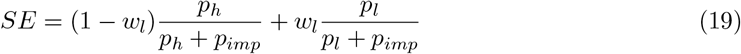

where *wl* and (1 − *wl*) are weights associated with pure lower and pure bins respectively.

The SE ranges from [0-1] depicting none to complete separation of functional state on binned representation surface. For real MD data, SE*<*0.4 is bad while separation starts to appear with SE around 0.5 and SE≥0.5 represents reasonable separation of functional states. SE serves as strict metric and SE=1 is practically impossible for real MD data.

*w_imp_* is a system specific parameter to reweigh the impure bins, as MD simulation data of different protein systems can have different levels of inherent impurity. For instance, intrinsically disordered protein have more inherent impurity in defining functional states as compared to folded proteins. *w_imp_* was assigned a value [0-1] such that supervised-RF has a maximum possible SE (with respect to no-classifier), hence it served as a strict test for URF. This is with the assumption that supervised-RF protocol works and no-classifier protocol does not. For systems like 32 bead LJ polymer, where no-classifier approach also works, no weights were provided (*w_imp_* = 0.5). Note that *w_imp_* does not weigh down the impure and can have values *>*0.5. *wl* is a data specific parameter as *X*^*MD*^ from MD simulations are generally imbalanced, such that a functional state can be a minor state (rare event) in terms of population. Hence, separation of pure lower and pure higher bins are weighted separately as per their population with *wl* = *population*(*functional state*).

### Functional regions

For T4L, the functional residues were directly visualized by depicting the input features (native contacts) as lines joining the residue pairs defining contact. The width of line represent mean (n=5) URF derived importance scores of top 200 features, same as used in representation plots. For mopR, the input feature scores for dihedrals were converted to residue level. For 5 cross validation iterations, if a residue was part of a dihedral with high URF importance score, the score of +1 was attributed to that residue. Final scores were normalized [0-1], depicted as width of spheres for each residue. For mopR, in addition to dihedral space of all residues, the URF was also trained on whitened coordinates of residue 20-220 in accordance with DiffNet (supplementary methods).

### Markov state models

The markov models were built with usual protocol ^48^ as implemented in pyemma library.^65^ The input *X*^*MD*^ is dimensionally reduced by TICA, discretized into *k* clusters by kmeans algorithm, time lagged transition matrix built at *ltm* lag time, and finally decomposed into *p* metastable pcca states. The *X*^*MD*^ is used as is for no-classifier approach, while the selected features were used for supervised-RF and URF integrated MSM (mopR: 40/229, *α*-synuclein: 200/9180). To account for hyper-parameter space, *k*=[20,30,50,70,100,150,250,350,500,700,1000,1500], *ltm*=(5,10..50) and (5,10..70) steps, and *p*=[3,4,5,6] and [2,3,4,5,6] were used for mopR and *α*-synuclein respectively. The incorrect hyper-parameter sets were detected if not all metastable states have a non-zero probability, or not all microstates were assigned to metastable states, or not all datapoints were used in pcca state decomposition, or convergence warning was raised. Five fold cross validations were performed for supervised-RF and URF integrated MSM. The ultimate output of MSM was metastable state labels (*L*^*p*^) corresponding to each instance of *X*^*MD*^.

Let say there exist a ground truth *L*^*T*^, a set of actual and manually defined state labels for *X*^*MD*^. For instance, the binding process in mopR is enrouted with unbound, bound and intermediate states ^35^ (supplementary methods, data-2). The identity (I) and purity (h) of MSM-derived metastable states (*L*^*p*^) can be defined in terms of ground truth *L*^*T*^ as:

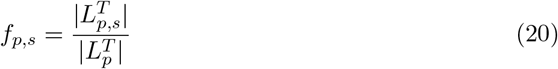

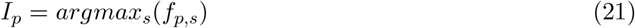

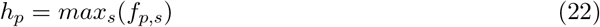

where 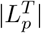 represents population of *L*^*T*^ belonging to *p*^*th*^ MSM derived metastable metastable state (*L*^*p*^), and 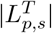 represents actual state population of *s*^*th*^ state in *L*^*T*^ .

Alternatively, purity of p-state markov model can be measured via weighted gini impurity guided by ground truth *L*^*T*^ as:

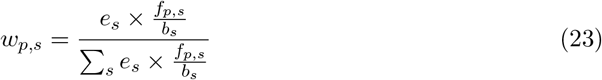

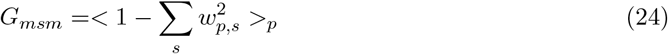

where, *e_s_* and *b_s_* are state probabilities corresponding to ideal distribution and *L*^*T*^ respectively.

In absence of ground truth, the goodness of MSM was measured as mean difference in all metastable state pairs in feature space by *fdiff* and in coordinate space by Davies Boulding index (*DBI*) ^66,67^ as:

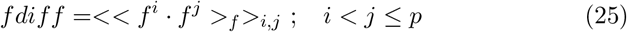

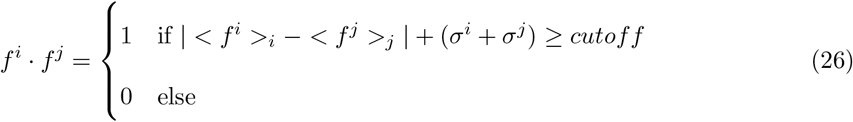

where *f* ^*k*^ is *f* ^*th*^ feature belonging to *k*^*th*^ metastable state, *σ* is standard deviation and *cutoff* = 0.51 was used.

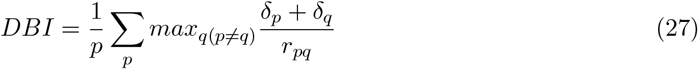

where, *δ_p_* represents average distance from data points of *p* metastable state to its centroid, while *r_pq_* is centroid distance between metastable states p and q.

### Baseline approaches

Important features were also selected as per VAMP2, SHAP and saliency scores, train-loo and predict-loo, RFE, amino, spectral oasis and MoSAIC protocols. Lower dimensional representations were also obtained by deepTICA, AE-TICA and TICA-VDE. Functional regions or conformational changes were detected by DiffNet. Metastable states corresponding to MSM derived states were identified using VampNets. Specific details are provided in supplementary methods.

## Supporting information

Supplemental methods, figure and results

## Code and data availability

All the codes and data used to generate the hereby reported results, the finalized URF implementation and tutorials are open sourced at https://github.com/msahilgit/Unsupervised-Random-Forest.

## Acknowledgement

We acknowledge support of the Department of Atomic Energy, Government of India, under Project Identification No. RTI 4007. All the authors acknowledge Tata Institute of Fundamental Research Hyderabad, India for providing the support of computing resources. JM acknowledges funding from Department of Science and Technology of India (CRG/2023/001426) MS would like to thank Palash Bera for helping in formalising mathematical details, Luigi Bonati for helping in deepTICA, Michael Ward for valuable suggestion on DiffNet, Sneha Menon and Subinoy Adhikari for assisting in IDP data and MSM analysis and Dheeraj Sarkar for his suggestions on imaging. MS also thanks Tejender Singh and Bhupendra Dandekar for valuable discussions.

## Supplementary Information

Extended Data Figs. 1-4, Supplementary Figs. 1-11, Supplementary methods and Supplementary table-1.

