## Supplemental methods, figure and results for "Unbiased learning of protein conformational representation via unsupervised random forest"

### Supplementary Information for Unbiased learning of protein conformational representation via unsupervised random forest

Mohammad Sahil,[*,†] Navjeet Ahalawat,[‡] and Jagannath Mondal[*,†]

†*Tata Institute of Fundamental Research Hyderabad, 36/P Gopanapalli Village, TS-500046, India*

‡*Department of Bioinformatics and Computational Biology, College of Biotechnology, CCS Haryana Agricultural University, Hisar 125 004 Haryana, India*

**Table of Contents**

| | |
|---|---|
| Extended Data Figs. 1-4 | 2-7 |
| Supplementary Figs. 1-9 | 8-20 |
| Supplementary Methods, Supplementary Figs. 10-11 | 21-35 |
| Supplementary Table 1 | 36 |

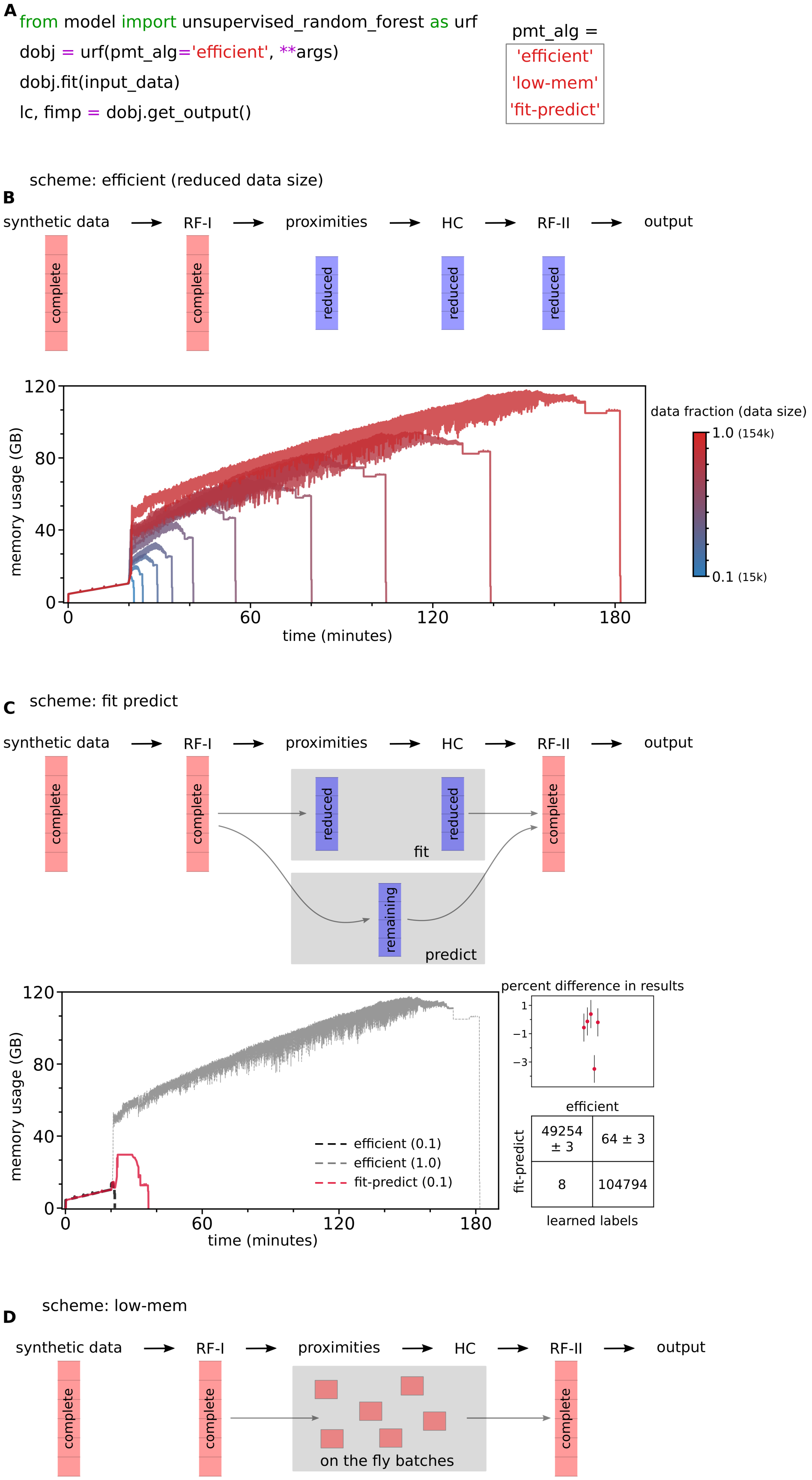
A
from model import unsupervised_random_forest as urf
dobj = urf(pmt_alg='efficient', **args)
dobj.fit(input_data)
lc, fimp = dobj.get_output()
pmt_alg =
'efficient'
'low-mem'
'fit-predict'
scheme: efficient (reduced data size)
B
synthetic data
RF-I
proximities
HC
RF-II
output
complete
complete
reduced
reduced
reduced
memory usage (GB)
time (minutes)
data fraction (data size)
1.0 (154k)
0.1 (15k)
C
scheme: fit predict
synthetic data
RF-I
proximities
HC
RF-II
output
complete
complete
reduced
reduced
complete
fit
remaining
predict
memory usage (GB)
time (minutes)
efficient (0.1)
efficient (1.0)
fit-predict (0.1)
percent difference in results
efficient
fit-predict
49254 ± 3
64 ± 3
8
104794
learned labels
D
scheme: low-mem
synthetic data
RF-I
proximities
HC
RF-II
output
complete
complete
on the fly batches
complete

**Extended Data Figure 1: Memory efficient implementation schemes for URF**: (A) A direct-user-friendly implementation of URF pipeline. (B) Time and memory usage for five fold RF-I and RF-II URF, with varying data fraction at memory intensive step via efficient scheme. (C) Time and memory usage for fit-predict scheme with 0.1:0.9 data in fit-predict part respectively, and its comparison with efficient scheme at 0.1 and 1.0 data size. The percent difference in results (SE) and confusion matrix for ($L^{hc}$) of fit-predict with respect to efficient (1.0) (n=5). (D) Schematic for low-mem scheme, which involve on-the-fly estimation of equation-3 without saving complete data, hence preventing memory overhead and performance but is very slow (few hours to days). Mostly efficient(1.0) scheme was used in this work. *Hardware specifications*: The benchmarking was performed on Rocky Linux 8.8 (Green Obsidian), AMD EPYC 7643 48-Core, 2.7-3.6 GHz, 96 cpus, 512 GB RAM machine.

---

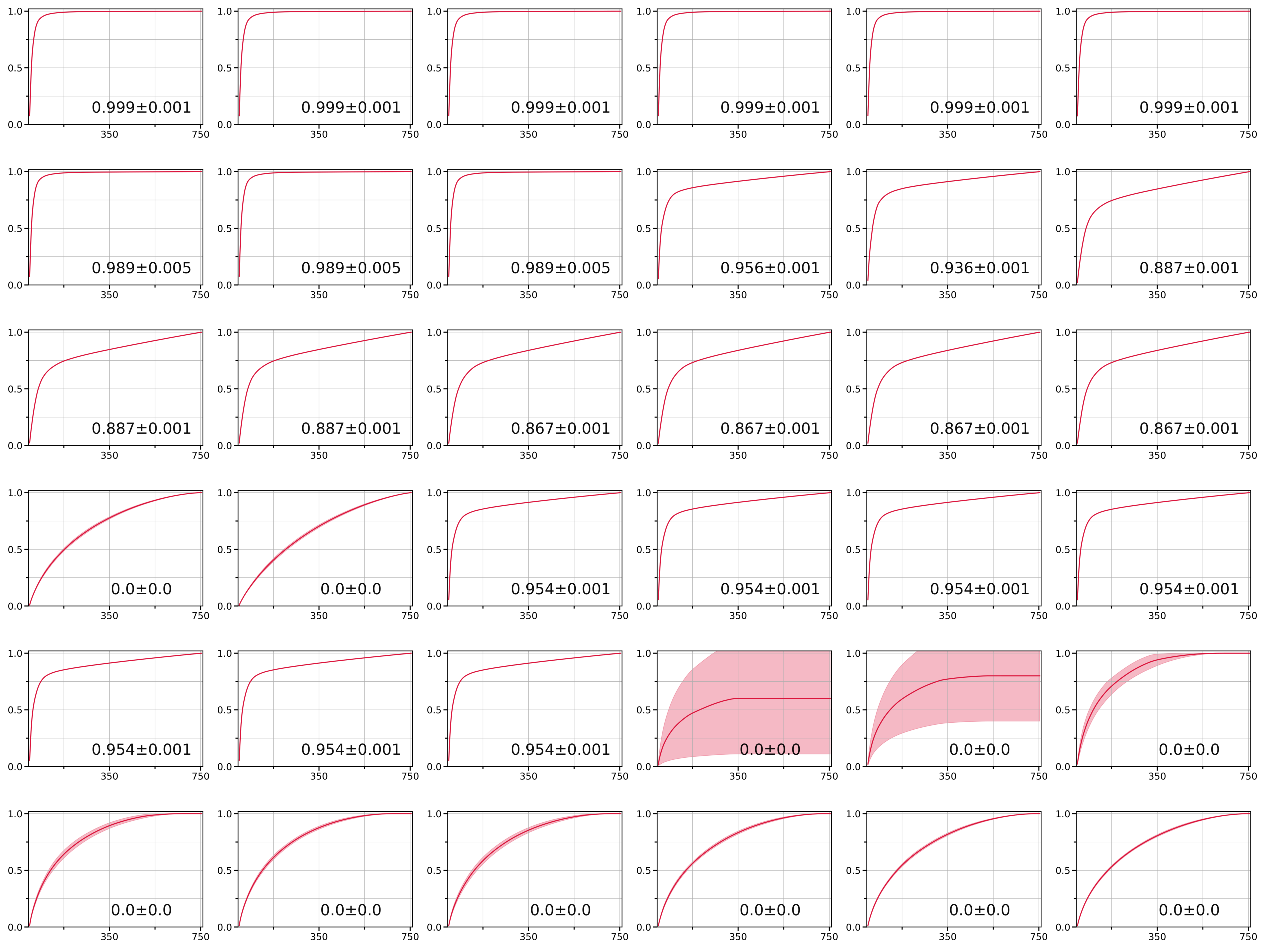


**Extended Data Figure 2: LC as predictive of importance curves**: The cumulative sum of URF derived importance scores increases almost vertically for initial top selected features, until an inflection point, followed by slow saturation to a total of 1. The curves from successfully trained URF (high LC values) have a higher inflection points and faster saturation to 1, compared to low LC URF models. The unsuccessfully trained URF (LC=0) exhibit almost zero vertical increase and/or large error bars. (x-axis = number of selected top features; y-axis = cumulative sum of URF derived feature importance scores; line and color represent mean and error (n=5)).

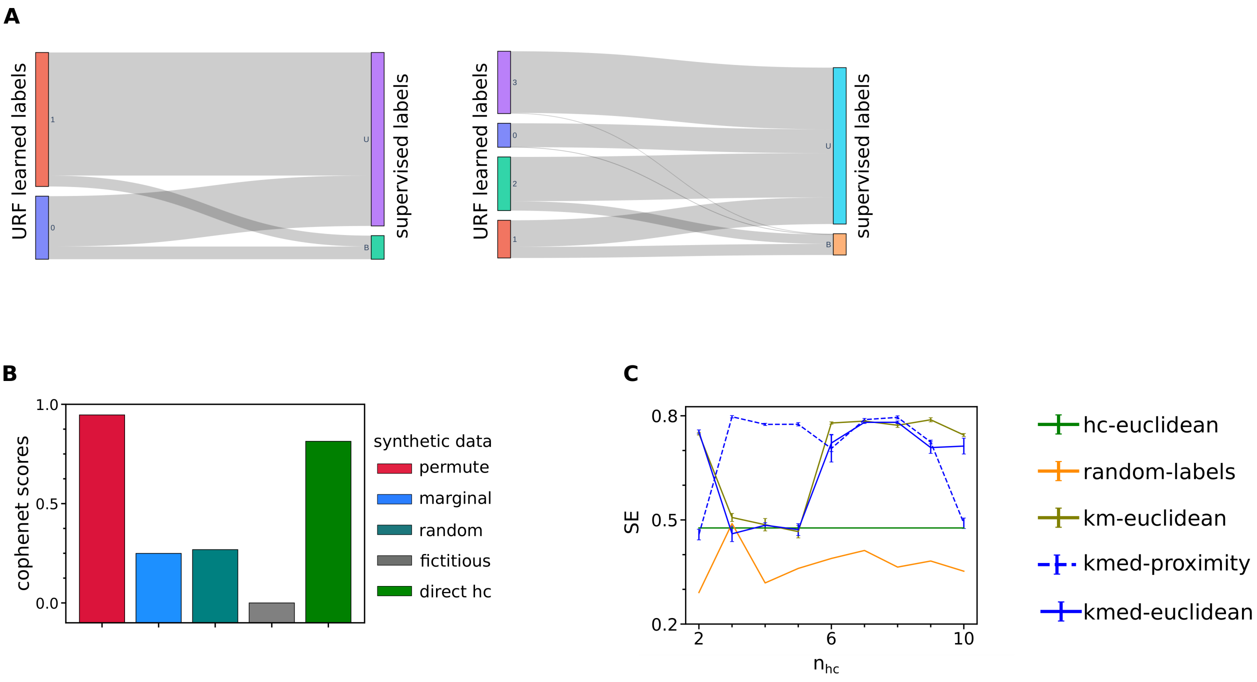


**Extended Data Figure 3: Independence of $n_{hc}$ in URF**: The URF performance can depend on $n_{hc}$ (number of hierarchical classes in data, main results), but it does not require apriori knowledge about system of interest, as $n_{hc}$ represent internal structure in data and does not corroborate actual labels. (A) The relation between URF learned $n_{hc}$ and actual labels (based on prior knowledge) via sankey plots, on T4L. (B) Cophenet scores, a metric for goodness of hc clustering, does not dictate URF performance. (C) Supervised-RF trained on labels derived from various clustering schemes yield unstable performance, potentially because these schemes are stable only at large n (number of clusters). hc, km and kmed represents hierarchical, kmeans, and kmediods. (n=5, errorbars for hc-euclidian and random-labels were large (0.265 and 0.07-0.16 respectively) and not shown for clarity.)

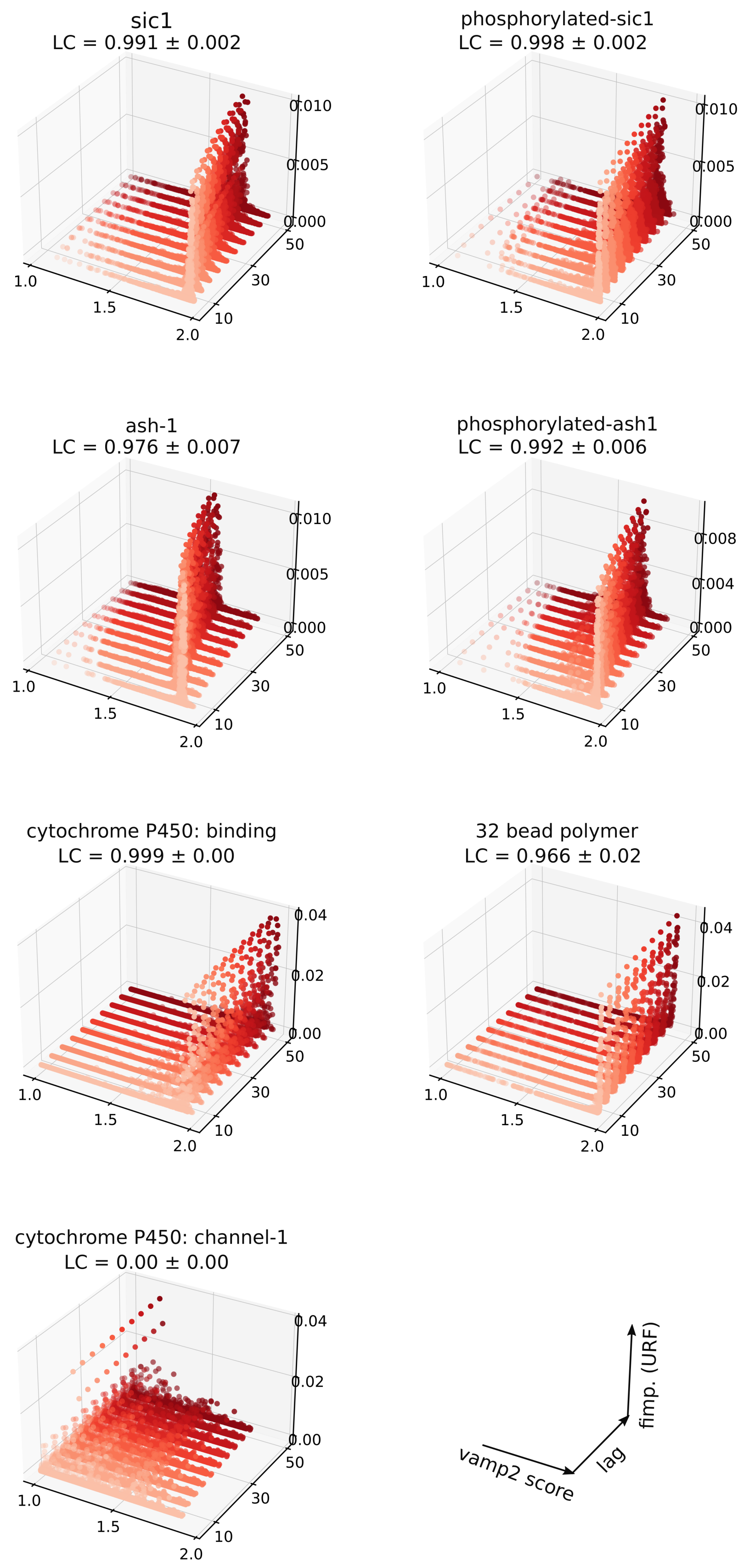
sic1
LC = 0.991 ± 0.002
phosphorylated-sic1
LC = 0.998 ± 0.002
ash-1
LC = 0.976 ± 0.007
phosphorylated-ash1
LC = 0.992 ± 0.006
cytochrome P450: binding
LC = 0.999 ± 0.00
32 bead polymer
LC = 0.966 ± 0.02
cytochrome P450: channel-1
LC = 0.00 ± 0.00
vamp2 score
lag
fimp. (URF)

**Extended Data Figure 4: LC predictive of fimp-VAMP2 relation**: The correlation between vamp2 scores and URF derived fimp scores, along with LC for different protein systems. For succesfully trained URF with high LC, the fimp-vamp2 relation is starkly evident, while not for cytochrome P450 (channel-1) with LC=0. For IDPs with faster dynamics, the fimp-vamp2 relation disappears at larger lag times as also seen for $\alpha$-synuclein (main). (The values represent mean, errorbars not shown for clarity).

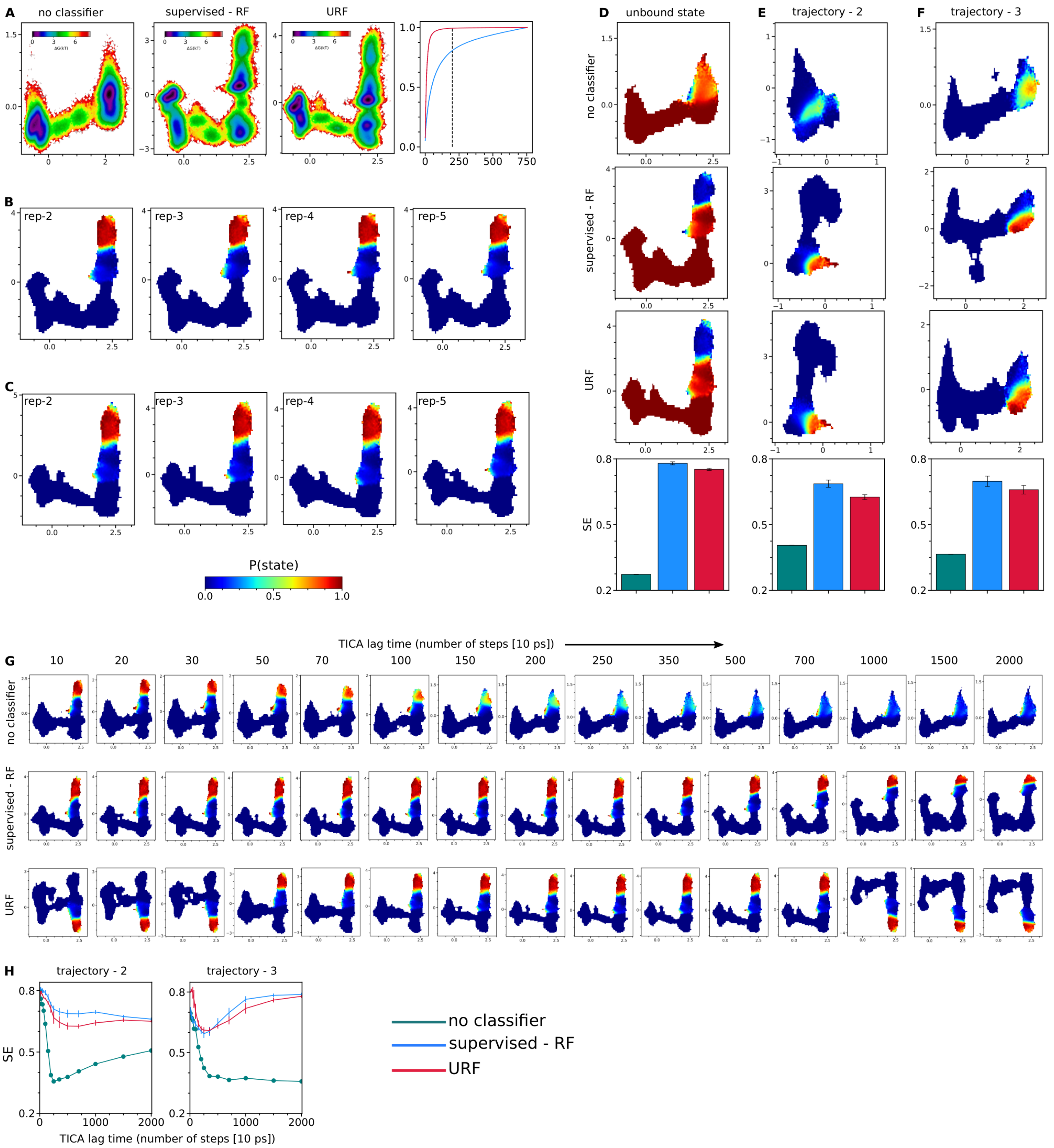


**Supplementary Figure 1:** (A) The TICA based free energy surface used in Fig. 2A and cumulative feature importance scores of supervised-RF and unsupervised-RF. (B, C) Cross validation replicates of bound state representation plots for supervised-RF and URF. (D,E,F) Representation plots for unbound state (traj-1) and bound state (traj-2,3) for T4L via no-classifier, supervised-RF and URF schemes and quantitative comparison via SE. (G,H) Representation plots at different TICA lag times (traj-1) and curves for traj-2, 3. (n=5).

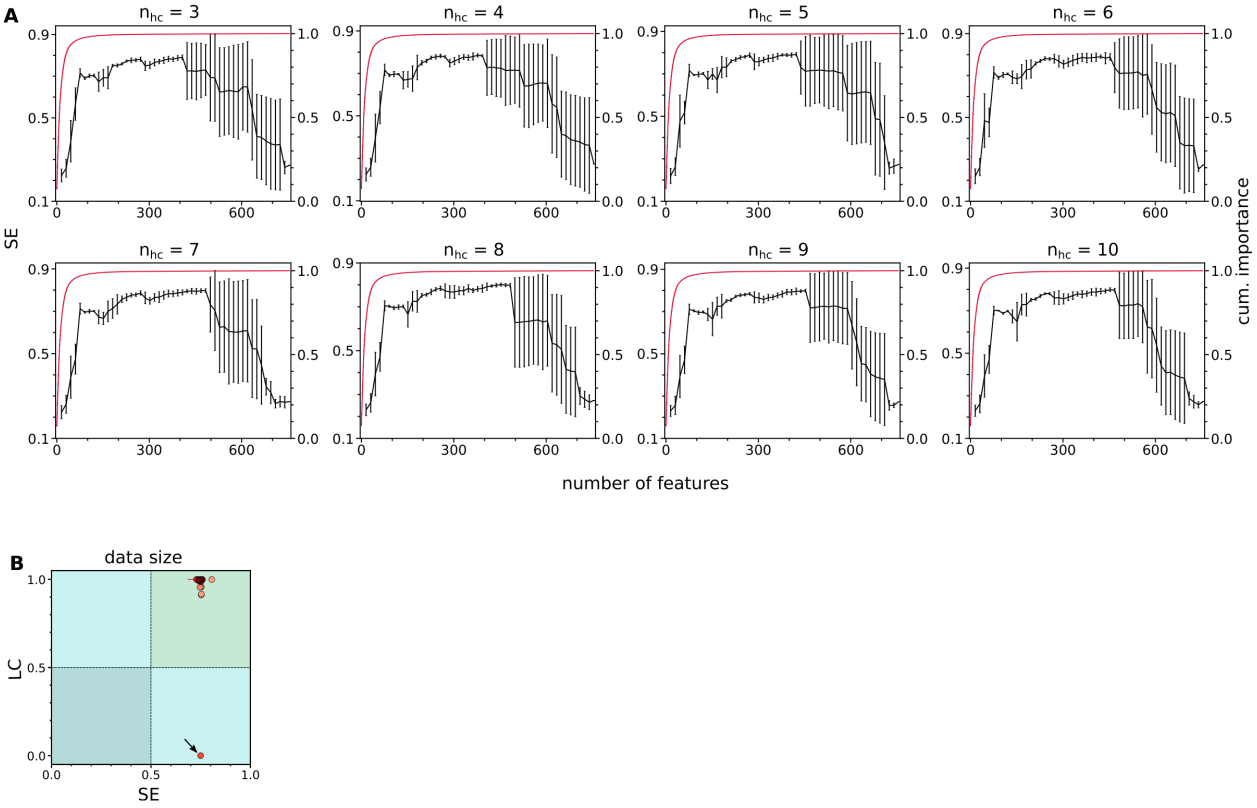


**Supplementary Figure 2:** (A) URF performance (SE) with increasing number of selected features for different $n_{hc}$, in extension to Fig. 2D ($n_{hc} = 2$). (B) Learning coefficient estimated for varying data size (efficient scheme, Fig. 2E) for different $n_{hc}$, correctly detects the URF performance except one case of false negative (arrow). The light to dark red color represent increasing $n_{hc}$. (n=5).

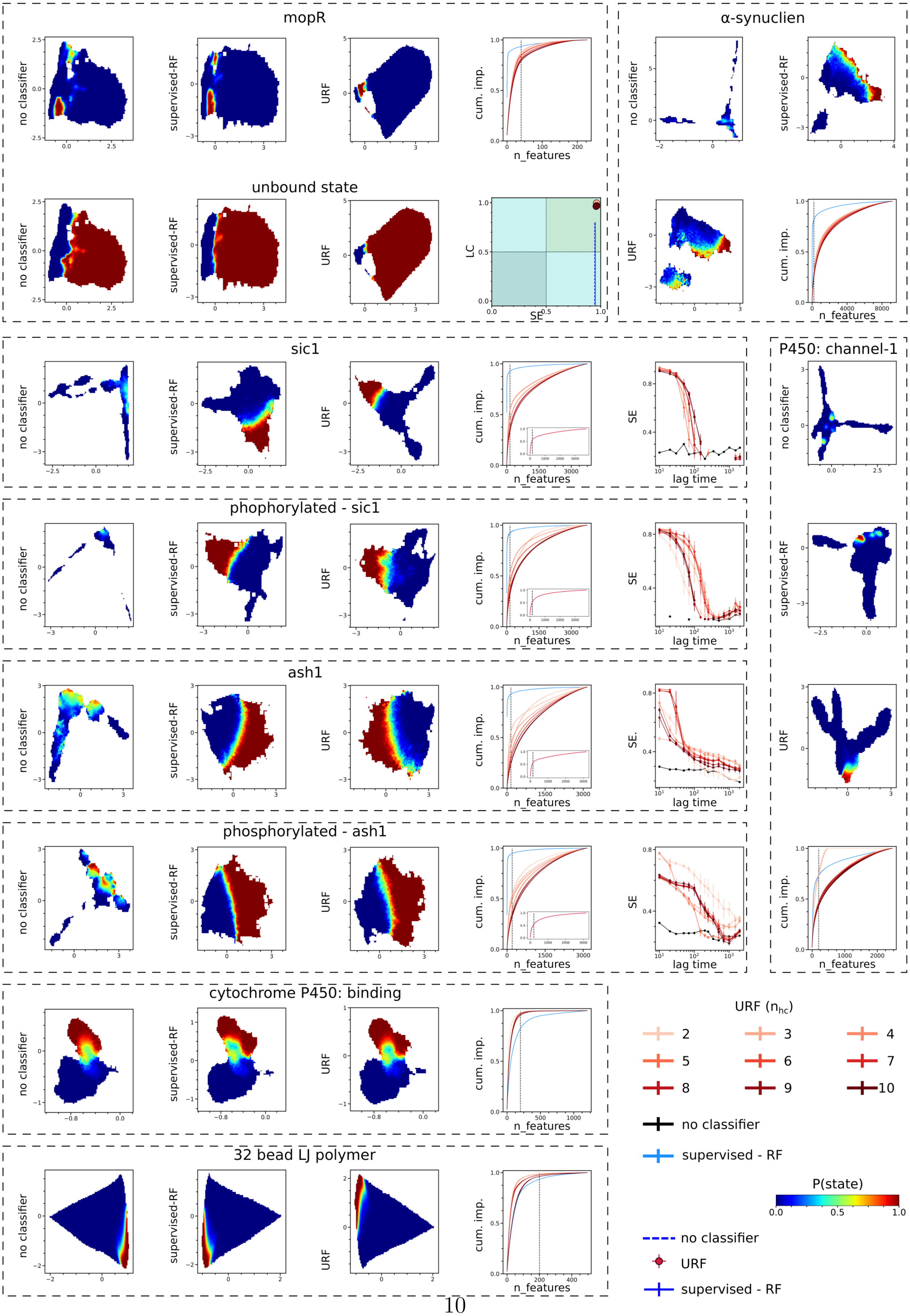
mopR
α-synuclien
unbound state
sic1
P450: channel-1
phophorylated - sic1
ash1
phosphorylated - ash1
cytochrome P450: binding
32 bead LJ polymer
no classifier
supervised-RF
URF
cum. imp.
n_features
LC
SE
lag time
URF ($n_{hc}$)
2
3
4
5
6
7
8
9
10
no classifier
supervised - RF
P(state)
no classifier
URF
supervised - RF

**Supplementary Figure 3:** Representation plots for different systems with no-classifier, supervised-RF and URF schemes. For a three-state system mopR, two states are shown (intermediate and unbound). Cumulative feature importance curves are shown for supervised-RF and URF ($n_{hc}$=2-10), with dashed line indicating the top features selected and width of curve as error bar (n=5). For intrinsically disordered proteins exhibiting faster dynamics, decrease in representation quality with increasing TICA lag time (number of time steps) is shown (n=5).

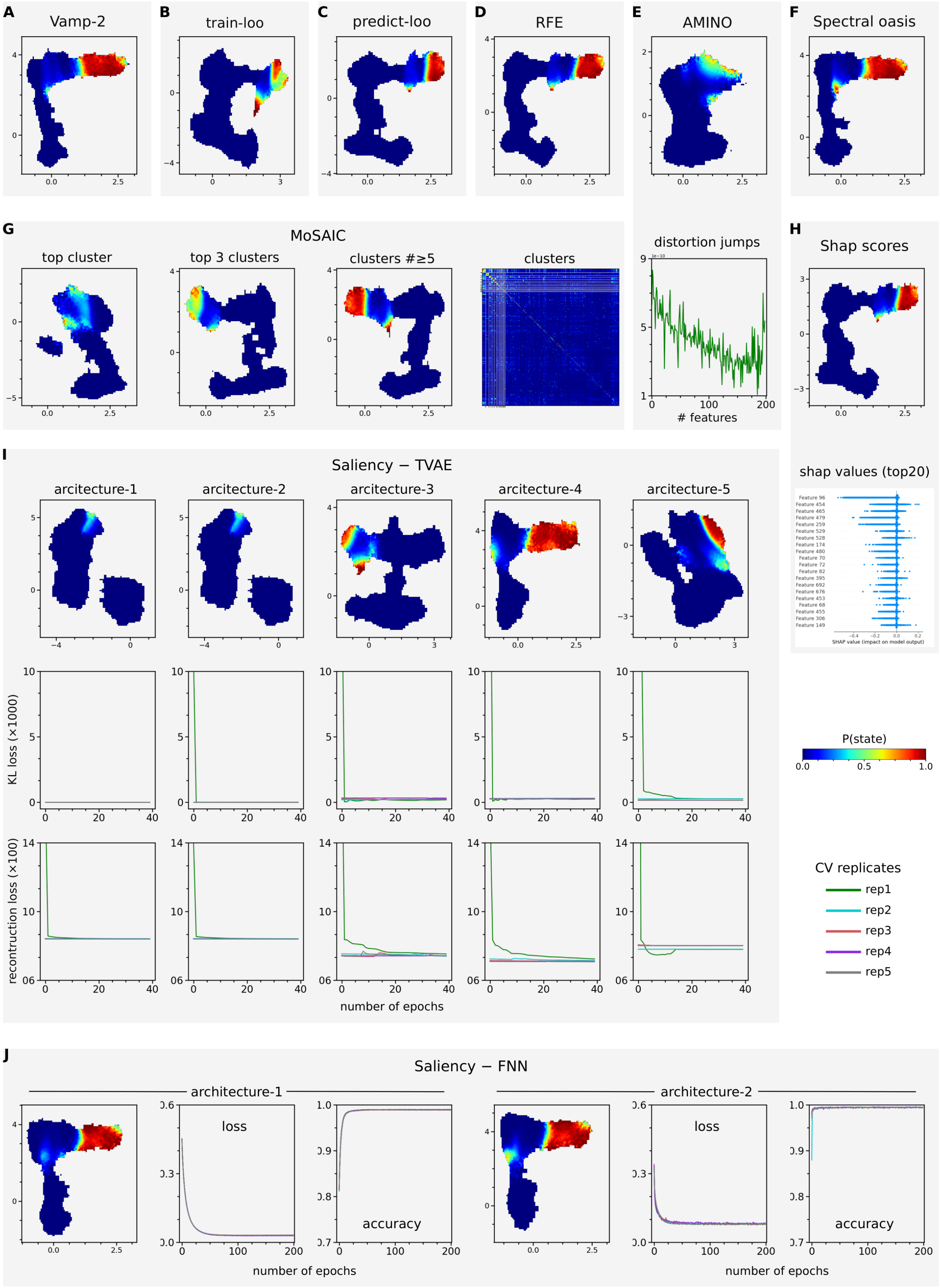
A Vamp-2
B train-loo
C predict-loo
D RFE
E AMINO
F Spectral oasis
G MoSAIC
top cluster
top 3 clusters
clusters #≥5
clusters
distortion jumps
### features
H Shap scores
shap values (top20)
SHAP value (impact on model output)
I Saliency − TVAE
arcitecture-1
arcitecture-2
arcitecture-3
arcitecture-4
arcitecture-5
KL loss (×1000)
recontruction loss (×100)
number of epochs
P(state)
CV replicates
rep1
rep2
rep3
rep4
rep5
J Saliency − FNN
architecture-1
architecture-2
loss
accuracy
number of epochs

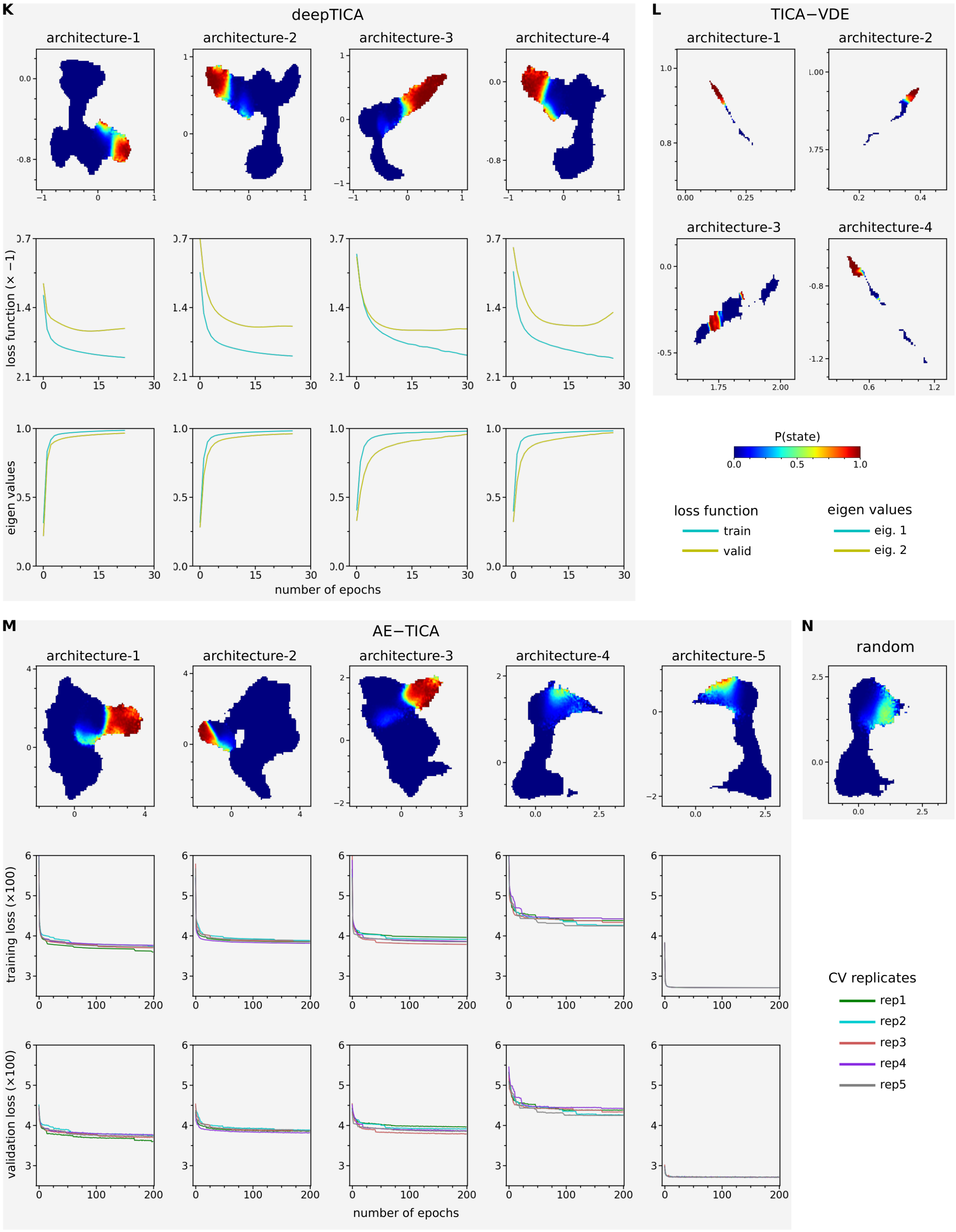


**Supplementary Figure 4:** Representation plots, learning curves and specific details of baseline approaches for T4 Lysozyme, corresponding to Fig. 2F (top plot).

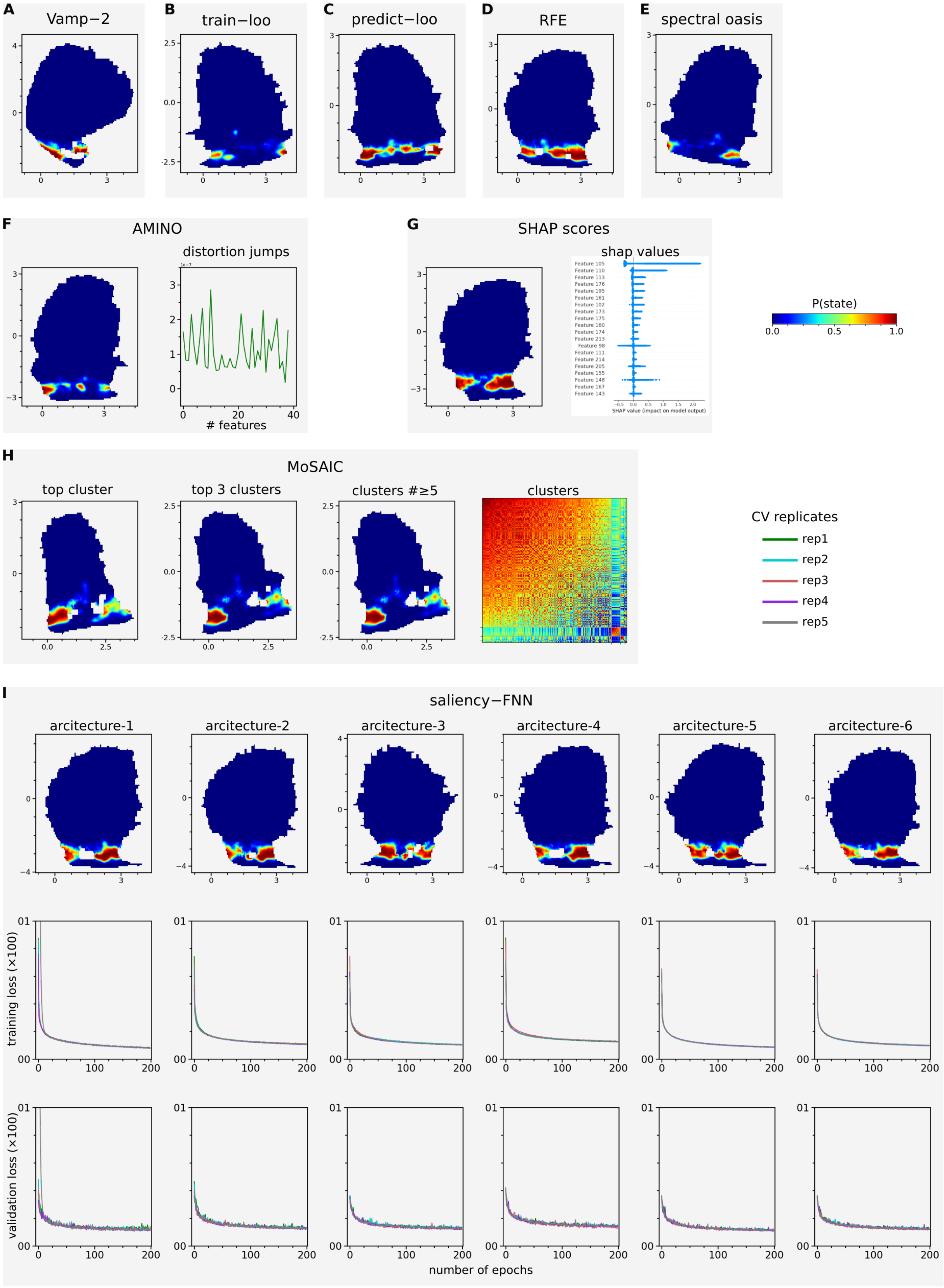
A
Vamp−2
B
train−loo
C
predict−loo
D
RFE
E
spectral oasis
F
AMINO
distortion jumps
### features
G
SHAP scores
shap values
SHAP value (impact on model output)
P(state)
0.0
0.5
1.0
H
MoSAIC
top cluster
top 3 clusters
clusters #≥5
clusters
CV replicates
rep1
rep2
rep3
rep4
rep5
I
saliency−FNN
arcitecture-1
arcitecture-2
arcitecture-3
arcitecture-4
arcitecture-5
arcitecture-6
training loss (×100)
validation loss (×100)
number of epochs

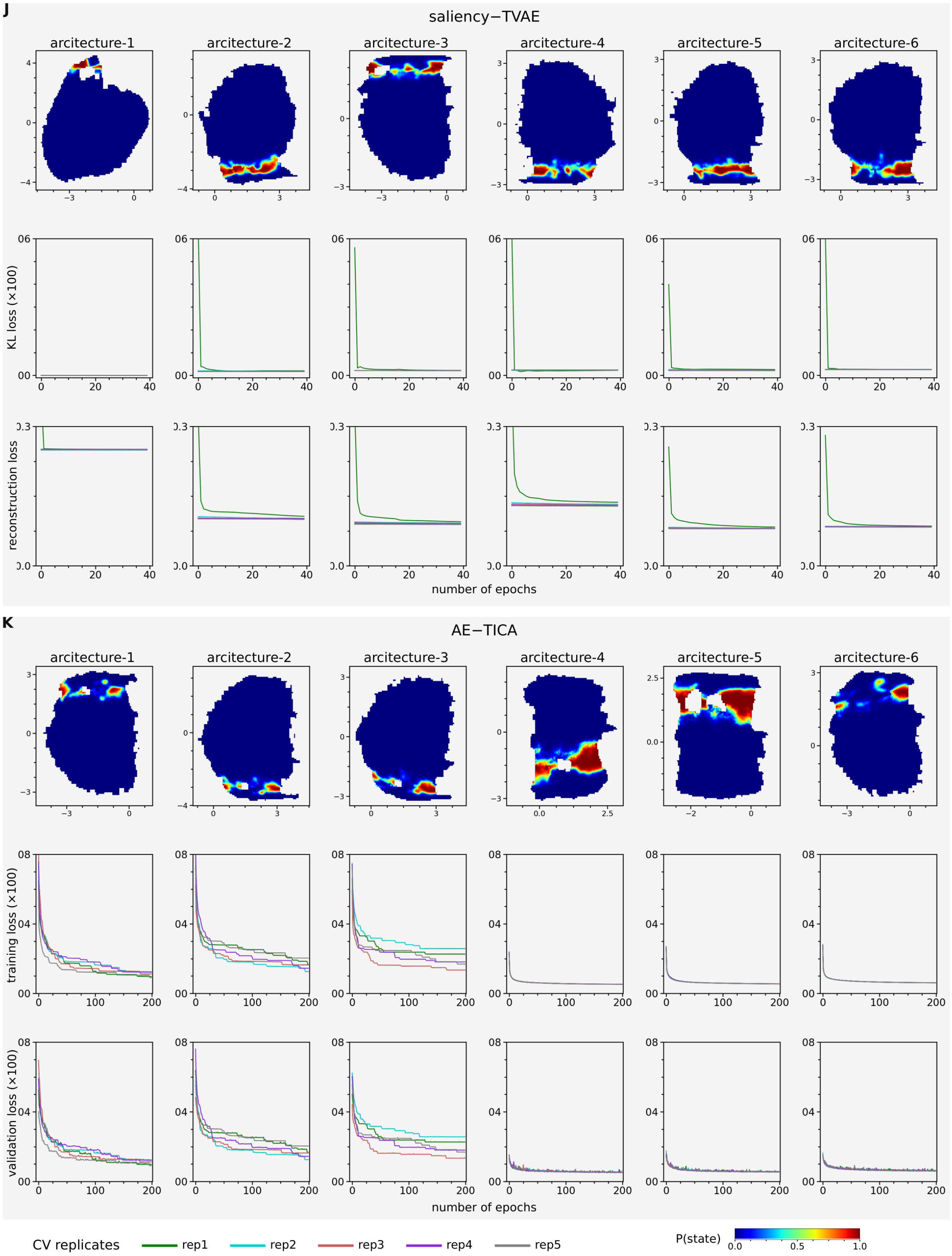
J
saliency−TVAE
arcitecture-1
arcitecture-2
arcitecture-3
arcitecture-4
arcitecture-5
arcitecture-6
KL loss (×100)
reconstruction loss
number of epochs
K
AE−TICA
arcitecture-1
arcitecture-2
arcitecture-3
arcitecture-4
arcitecture-5
arcitecture-6
training loss (×100)
validation loss (×100)
number of epochs
CV replicates
rep1
rep2
rep3
rep4
rep5
P(state)
0.0
0.5
1.0

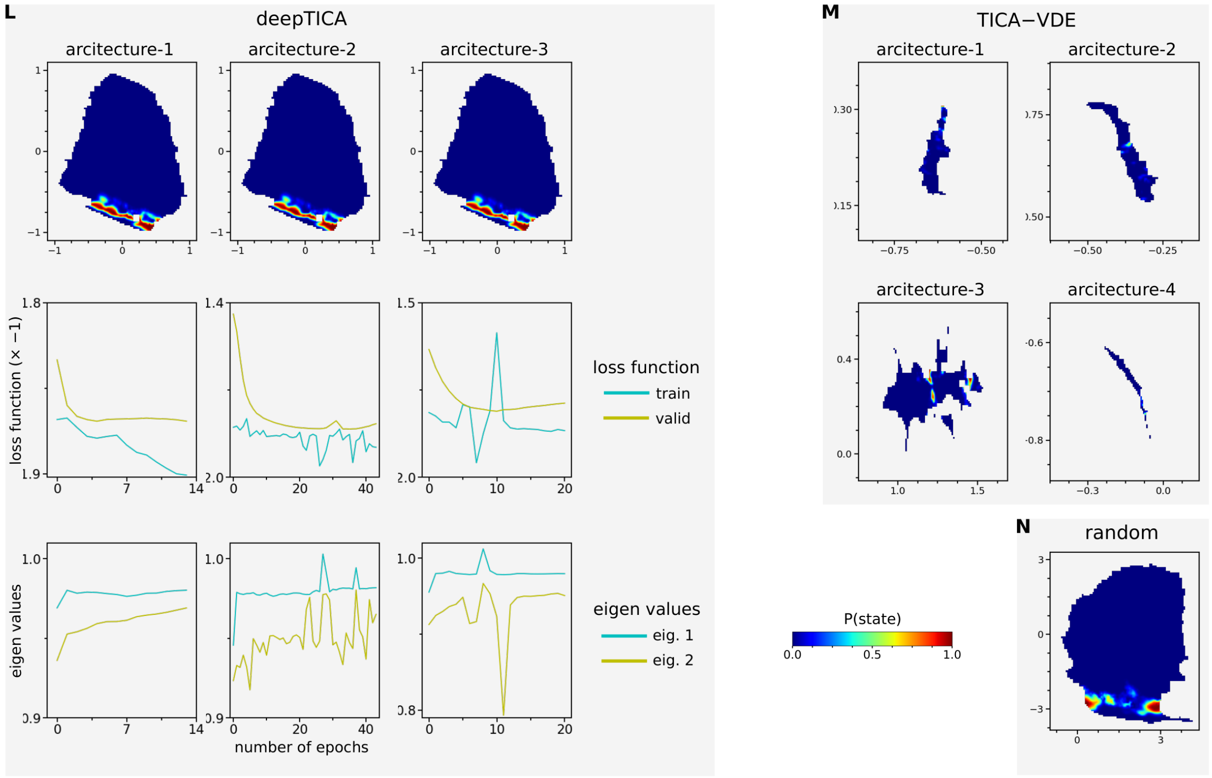


**Supplementary Figure 5:** Representation plots, learning curves and specific details of baseline approaches for mopR, corrosponding to Fig. 2F (bottom plot).

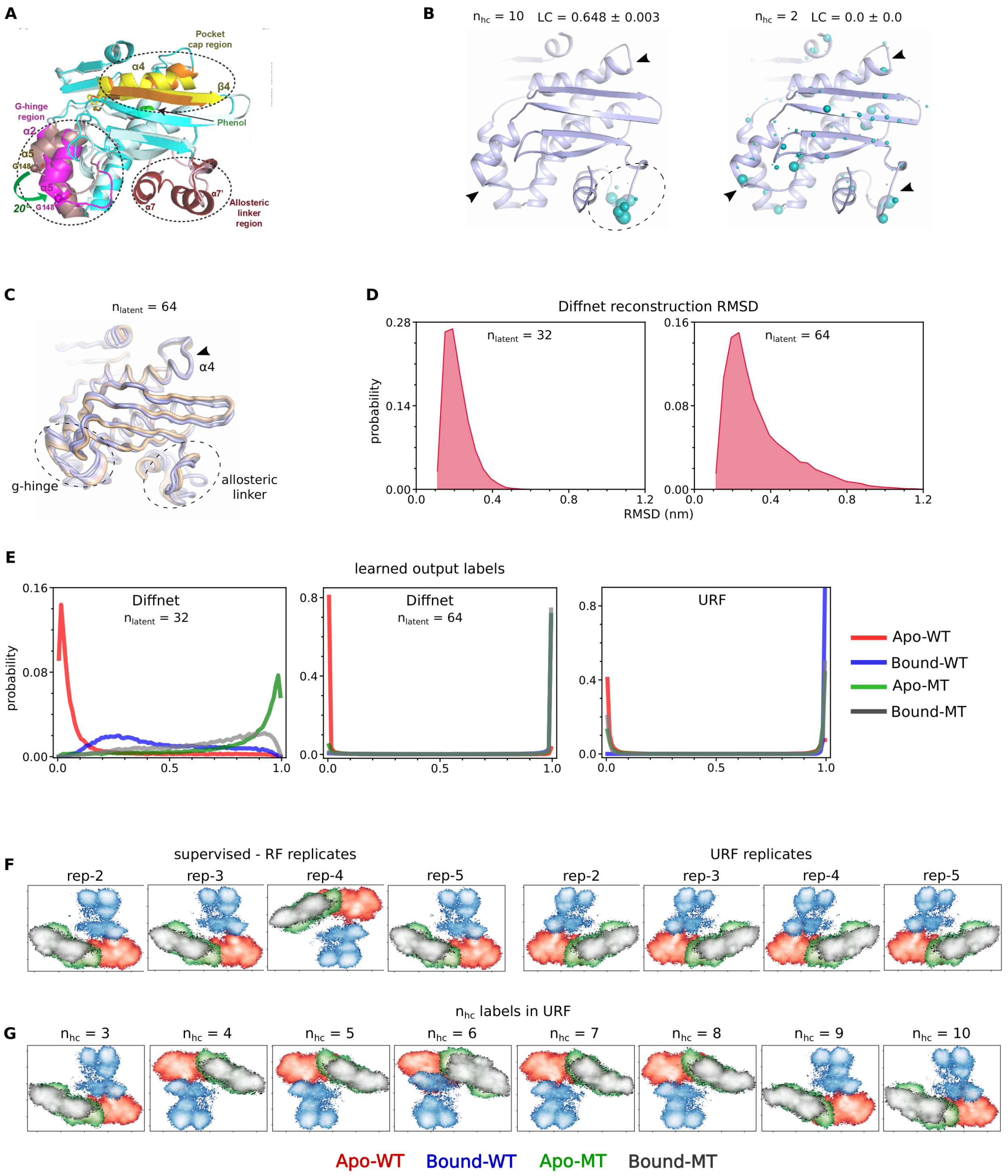


**Supplementary Figure 6:** (A) The three functional regions of mopR identified via simulations based hydrogen bonding network and mutagenesis based tryptophan quenching studies. G-hinge region not visible in crystallography is emphasized and $\alpha 4$ region is part of pocket cap region. (copyright here) (B) Selected residues of mopR as identified by partially successful and unsuccessful URF models (as per LC values). These URF models were trained on whitened coordinates. (C) Functional regions identified by DiffNet models trained with $n_{latent} = 64$ as opposed to $n_{latent} = 32$ in maintext. The dashed circles and arrowheads in (B) and (C) indicate successfully identified and missed functional regions respectively. (D) The root mean squared deviations of ...

... DiffNet reconstructed structures with respect to actual. (E) The learned output labels of DiffNet and URF for different states of mopR. (F, G) FES representation plots of mopR states via cross validation of supervised-RF or URF ($n_{hc} = 2$) and of URF with varying $n_{hc}$.

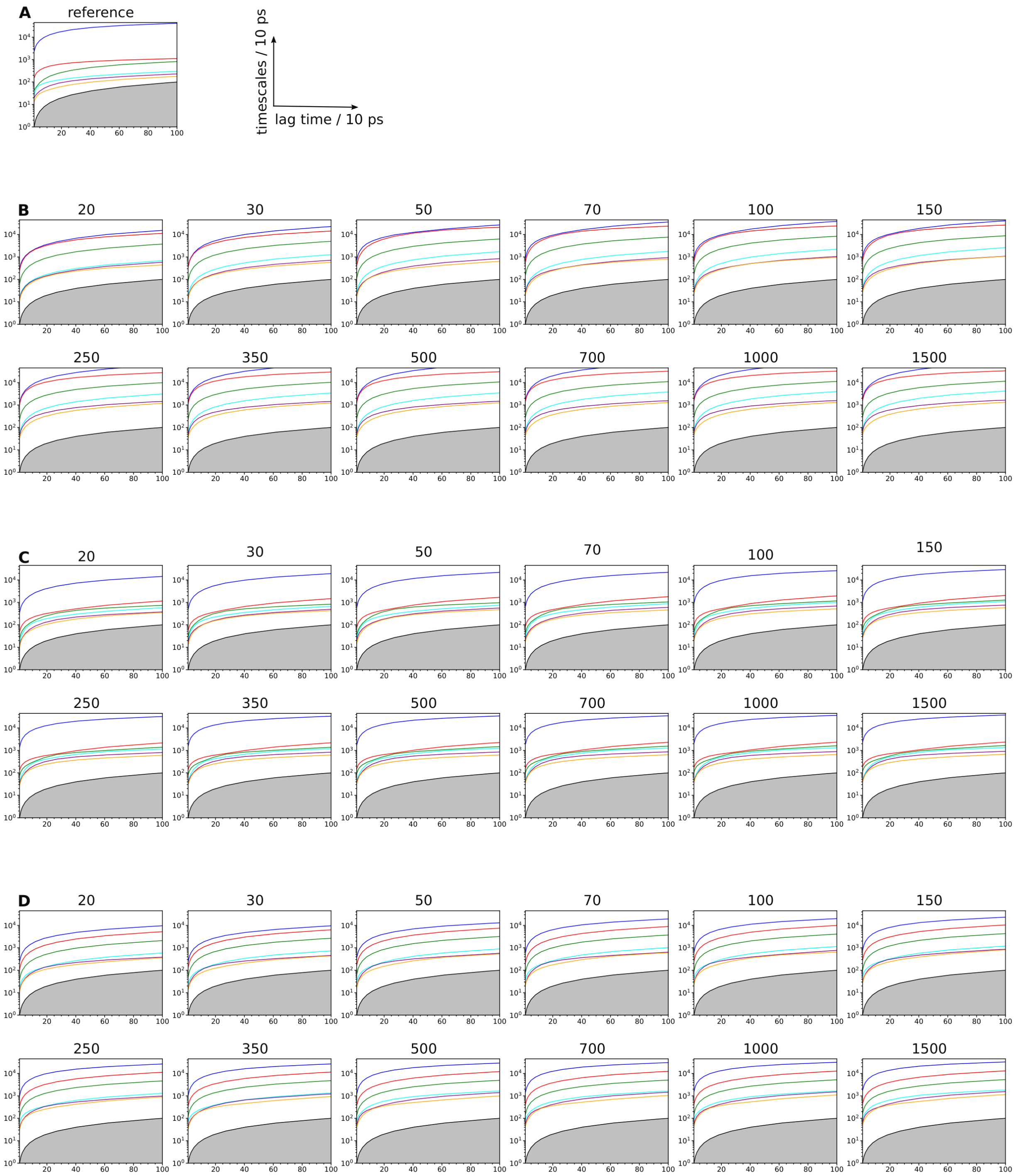


**Supplementary Figure 7:** The implied timescale plots of reference msm[1] (A), no-classifier (B), URF (C) and supervised-RF(D) at varying number of clusters.

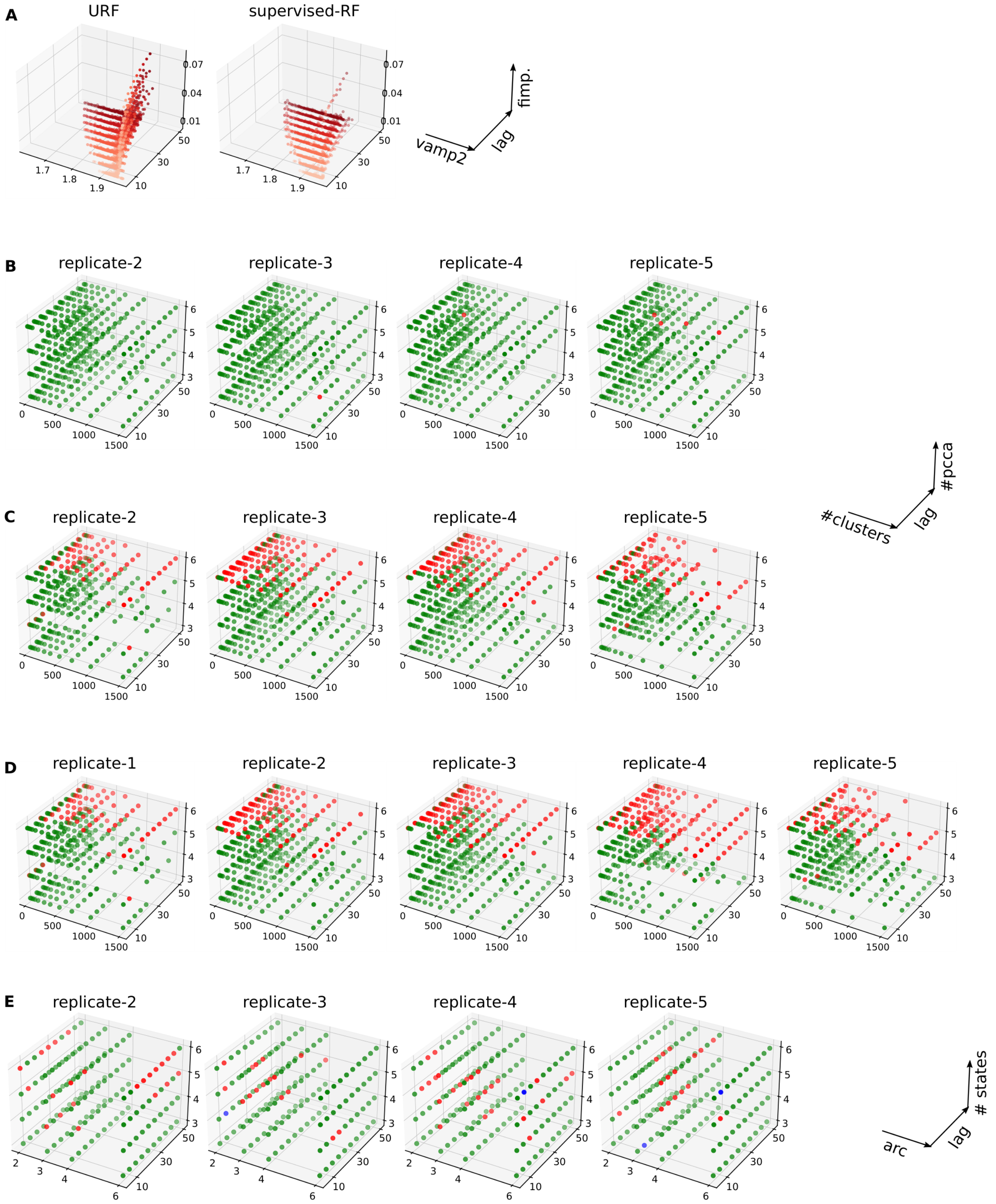


**Supplementary Figure 8:** (A) Vamp-2 fimp relation for URF ($n_{hc} = 3$) and supervised-RF. (B-E) MSM derived via cross-validation replicates of supervised-RF (B), URF ($n_{hc} = 2$) (C), URF ($n_{hc} = 3$)(D) and vampnet (E).

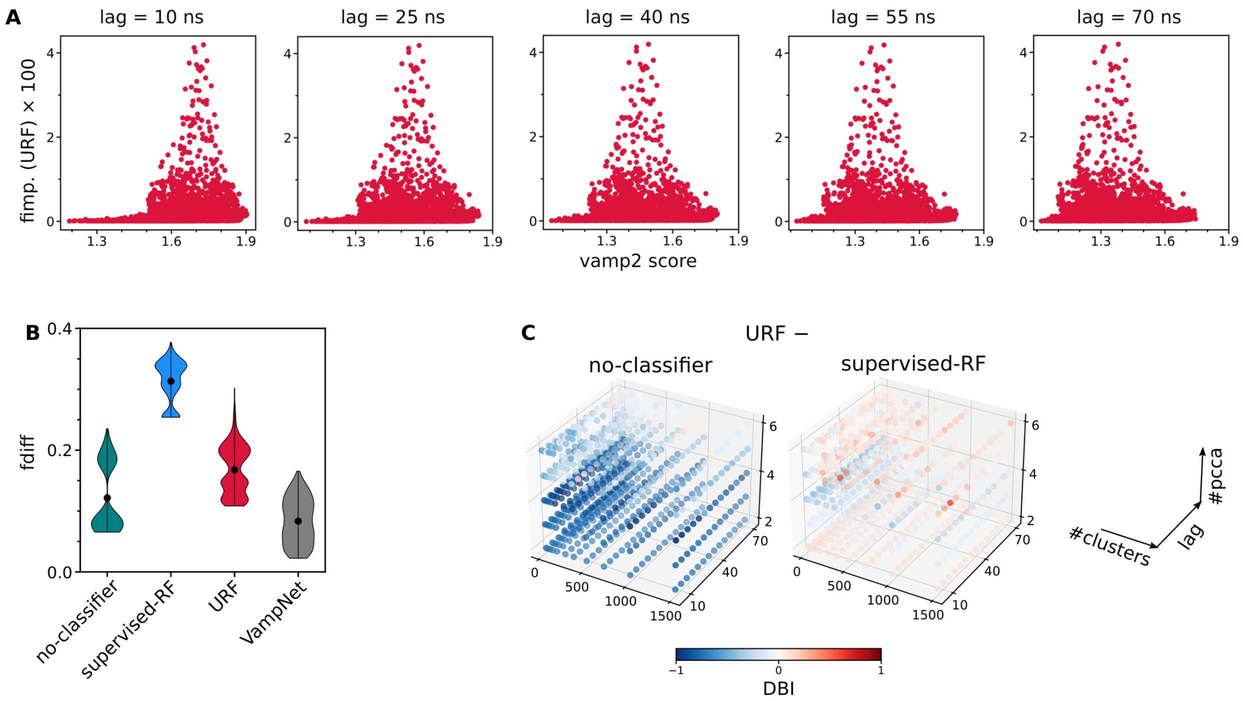


**Supplementary Figure 9:** (A) The vamp2-fimp(URF) relation for $\alpha$-synuclien at increasing lag times. (B) The ensemble comparison of *fdiff* values estimated for markov models via different schemes, in addition Fig. 4I. (C) One-to-one comparison of URF integrated msm with no-classifier and supervised-RF integrated msm via *DBI*. (The values represent mean (n=5), errorbars not shown for clarity.)

### Supplementary Methods

The protein systems and datasets used in this work are briefly described here. Specific, technical and complete details can be found elsewhere in respective references. For each system, the functional state and metrics to detect them were defined. For instance, the compact and extended conformations of intrinsically disordered proteins were respectively defined based on $r_g$ values lower or higher than experimentally observed $r_g$ value. These metrics were used to define the labels (continuous or categorical) for functional states used in supervised-RF training, measurement of SE, or defining reference states in markov models. These metrics were also used to train supervised baseline approaches like saliency-FNN, SHAP.

#### Data 1: ligand binding simulations of T4L

T4L is a lysozyme from the T4 bacteriophage. Its binding site mutant L99A with increased 150 $\AA^3$ pocket volume represent a model system for protein-ligand binding studies. Unbiased binding simulations of benzene to L99A-T4L were used from previous work.[2] These simulations were started with protein and multiple ligand copies (2 benzene molecules) in unbound state. Ligand molecules were unbiasedly allowed to find its path to binding pocket, ultimately resulting in binding i.e., crystallographic bound pose. Total of six independent simulations were used amounting to 27.4 $\mu$s of data saved at an interval of 10 ps i.e., 2741970 datapoints. Native contacts were defined as residue pairs with $C_\alpha - C_\alpha$ distance within 10 $\AA$ as measured in crystallographic structure (pdb-3DMX). These 755 dimensional native contacts define protein conformation at any instant of time and was correlated with bound and unbound state of ligand. $w_{imp}$ was found to be 0.44 for trajectory 1, mainly used in this work. Bound and unbound states were defined based 0.6 nm cutoff from binding pocket, as defined previously.[3] Bound state was used for detecting conformational separation of functional states, with $w_l = 0.066$ i.e., population fraction of bound state in trajectory 1. $w_{imp} = 0.44$ was estimated to be 0.44 for trajectory-1.

#### Data 2: ligand binding simulations of MopR

MopR is a bacterial enhancer binding protein which activate catabolic genes for phenol degradation in *Acinetobacter calcoaceticus*.[4] Its N-terminal domain acts as phenol sensor and is a potential candidate for phenol-pollutant biosensor.[5] Its N-terminal domain(residues 1-229) was used for performing unbiased ligand binding simulations.[1] Dimeric mopR with 1:5 protein:ligand (aromatic pollutant phenol) concentration in unbound state were simulated to unbiasedly yield a crystallographic bound pose (pdb id: 5KBE) in any one or both protomer. Total of 7 binding trajectories were generated. On top of these long and complete binding trajectories, 122 adoptive sampling simulations each of 100 ns were also generated. A total of 129 simulations were used amounting to around 29 $\mu$s, saved at an interval of 10 ps i.e., 2897865 data points. Protein-ligand contact map were generated i.e., distance between CZ atom of phenol and each CA atom of protein were calculated, giving a total of 458 dimensions. Since the two protomers of mopR dimer are identical, 458 dimensions were reduced to 229 by taking minimum of identical residue distances. Bound, intermediate and unbound states were defined as previously,[1] based on center of geometry distance between binding pocket and ligand with 0-0.35 nm as bound, 0.35-1 nm as intermediate and >1 nm as unbound states. $w_{imp}$ was estimated to be 0.4. $w_l$ was 0.699, 0.053 and 0.248 for bound, intermediate and unbound states respectively.

#### Data 3-6: Simulation ensembles of MopR

MopR has also been studied for its dynamic allostery, change in function without observable conformational change (pdb ids: 5KBE and 7VQF).[6] Its G148P mutant, far away from binding site, can increase the binding 7-fold ($k_d$, from 0.48±0.06 to 0.07±0.02 $\mu$M). Based on ligand dependent allostery and site directed mutagenesis, mopR was studied in 4 different states in our previous work:[6]

i. Apo state: wildtype without ligand, referred as apo-wt (data 3)

ii. Bound state: withtype with ligand bound in both monomers, referred as bound-wt (data 4)

iii. G148P apo: a distal mutant (affecting $k_d$ by 10x) but without ligand, referred as apo-mt (data 5)

iv. G148P bound: a distal mutant with ligand bound in both protomers, referred as bound-mt (data 6)

Simulations of dimeric mopR in each state were performed starting from bound-wt crystal structure (pdb-5KBE), for which other states were remodelled. Simulations were performed for atleast 1 $\mu$s (1.5 $\mu$s for bound-wt). A total of 10 simulations (apo-wt:3, bound-wt:1, apo-mt:3, bound-mt:3) were used, amounting to 10.5 $\mu$s saved at 20 ps i.e., [150001, 75000, 150001, 150001] datapoints. A 648 dimensional dihedral space was estimated consisted of $\phi$, $\psi$ and $\chi 1$ dihedrals for protomer1 residues. Note that for some residues like proline and glycine, some dihedrals were not possible. The four states served as labels for the respective simulations. In addition to dihedrals, the xyz coordinates of $N$, $C_\alpha$, $C_\beta$, and $C$ atoms corresponding to residues 20-220 were recorded for the aligned trajectories. The coordinates ($X \in \mathbb{R}^{(n,f)}$) were whitened as:

$$\overline{X} = X - < X >_n \tag{28}$$

$$C_{00} = \frac{1}{n-1}\overline{X}^T\overline{X} \tag{29}$$

$$\lambda, \nu = eig(C_{00}) \tag{30}$$

$$C_{00}^{-\frac{1}{2}} = \nu \frac{1}{\sqrt{\lambda}} \nu^T \tag{31}$$

$$X^{MD} = \overline{X} C_{00}^{-\frac{1}{2}} \tag{32}$$

where $\lambda$ and $\nu$ represent eigenvalue (diagonal) and eigenvector matrices. The final $X^{MD}$ was used to train URF. For mopR, the input data corresponding to all residues was used for training URF, but analysis was only considered for residues 20-220, for comparison with DiffNet.

#### Data 7,8: simulation ensembles of sic1 with and without phosphorylation

Sic1 is a cell-cycle regulator in *Saccharomyces cerevisiae* which is regulated by multiple phophorylations in its 92 residue long intrinsically disordered region. Its unbiased MD simulations with and without multiple phosphorylations were used in this work.[7,8] The unphosphorylated sic1 simulation (data 7) is a 30 $\mu$s long single trajectory, saved at a 180.036 ps time interval or 159729 datapoints.[7] The phosphorylated sic1 simulations (data 8) were performed with phosphorylations at Thr7, Thr35, Thr47, Ser71, Ser78 and Ser82. 25 independent simulations each of 500 ns were performed, amounting to 124900 datapoints saved at 100 ps time interval.[8] Simulations were treated as single trajectory. 3741 dimensional protein contact map was estimated as minimum distances between all residue pairs except neighbouring three residues. $w_{imp}$ was estimated to be 0.26 and 0.5 for sic1 and phosphorylated-sic1 respectively. The functional states were defined on the basis of $r_g$ (radius of gyration), such that states above and below experimentally observed $r_g$ were referred to expanded and compact states respectively. The experimental $r_g$s were 2.837 and 2.993 nm corresponding to $w_l$ of 0.161 and 0.159 in sic1 and phosphorylated sic1 respectively.

#### Data 9,10: simulation ensembles of ash1 with and without phosphorylation

83 residue long intrinsically disordered ash1, responsible for switching mating type and pseudohyphal growth in *Saccharomyces cerevisiae*, was simulated with and without phosphorylations.[7,8] The unphosphorylated ash1 (data 9) simulation is a 30 $\mu$s long single trajectory, saved at 180.036 ps time interval or 136050 datapoints.[7] Phosphorylations were introduced at Ser7, Ser9, Thr12, Ser25, Thr33, Ser35, Ser38, Ser48, Ser52 and Ser73. 25 independent and concatenated simulations each of 500 ns saved at 100 ps amounted to 124672 datapoints were utilized in this work.[8] 3160 dimensional contact map was estimated as for sic1. Functional states were also defined based on experimental $r_g$ of 2.85 and 2.75 nm for sic1 and phosphorylated sic1 corresponding to $w_{imp}$ of 0.57 and 0.70 and $w_l$ of 0.35 and 0.4 respectively.

#### Data 11: Simulation ensemble of $\alpha$-synuclein

$\alpha$-synuclein is a 140 residue long intrinsically disordered protein involved in Parkinson's disease via its aggregated insoluble fibrils and Lewy bodies. A single 73 $\mu$s long simulation trajectory was performed by D. E. Shaw Research, saved at 1 ns time interval amounting to 73124 datapoints.[9,10] A 9180 dimensional protein contact map was estimated as $C_\alpha - C_\alpha$ distances between all residue pairs expect nearest 5 neighbours. Extended state was defined based on $r_g > 3$ nm, corresponding to $w_{imp}$ of 0.23 and $w_l$ of 0.153.

#### Data 12: Open-closed simulation ensembles of cytochrome P450cam

An archetypal cytochrome P450, CYP101A1 - popularly known as P450cam,[11] constituted of substrate ingress channel-1 which can be in open and closed conformations. 9 independent simulations of substrate-free P450cam (starting structure-pdb id 3L61) sampling open and closed conformational extremes were utilized in this work.[12] Simulations were 500 ns to 1000 ns long, saved at 10 ps time interval amounting to 573009 datapoints. A 2461 dimensional native contacts were estimated to detect channel-1 associated conformational changes in P450cam, measured via residue-residue minimum distances for all residues having $C_\alpha - C_\alpha$ distance of $\leq 10\mathring{A}$ in starting structure. Open and closed states were separately defined as previously[12] based on RMSD values (with respect to closed structure) of F, G, F-G loop and $\beta'$ helices, such that rmsd within 2 $\mathring{A}$ defined the closed state, corresponding to $w_{imp}$ and $w_l$ of 0.19 and 0.10 respectively.

#### Data 13: binding simulations of P450cam

Unbiased binding simulations of P450cam with its cognate ligand camphor were utilized in this work.[13] Three independent simulations starting from unbound state of P450cam with 4 ligands in bulk solvent were performed unbiased until the binding in heme active site was achieved. Three simulations corresponds to 1274375 datapoints saved at 10 ps time interval (12.7 $\mu$s). 1203 dimensional native contacts were measured to define the protein conformations, as residue-residue minimum distances between all residue pairs with $C_\alpha - C_\alpha$ distance $\leq 10\mathring{A}$ in crystal structure 2CPP (pdb id). The functional states (bound state) was defined based on ligand's center of mass distance from heme moiety of $\leq 0.6$ nm.[3] The $w_{imp}$ was not weighted for this system, hence 0.5. The $w_l$ was found to be 0.116 for trajectory 1.

#### Data 14: Simulations of 32 beads LJ polymer

32 bead LJ polymer is a simple protein-like system consisting of linearly bonded carbon like atoms having only Lennard-Jones (LJ) interaction potentials. This system has served as a toy system for

studying multiple aspects of proteins.[14] In this work, we have utilized 200 independent simulations each of maximum 30 ns, starting from different extended to collapsed state. The trajectories were saved at 1 ps time interval amounting to 5978731 datapoints. A 496 dimensional contact map was estimated to define its conformation as used previously, measured as distances between all pairs of beads. The functional states were defined based on $r_g < 0.5$ nm (collapsed state), corresponding to $w_l$=0.16 and $w_{imp}$ unweighted at 0.5.

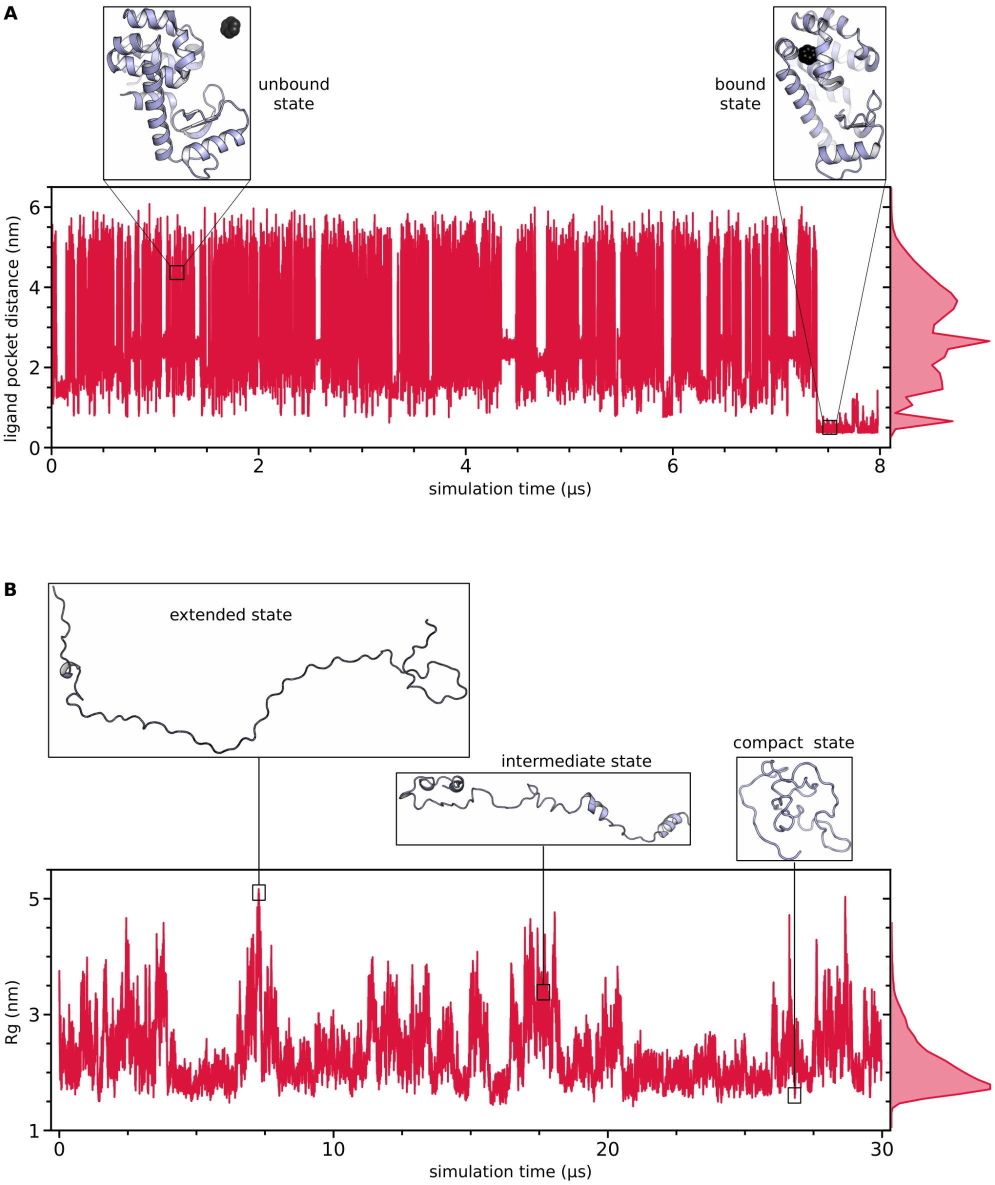


**Supplementary Figure 10:** (A) A typical unbiased ligand binding trajectory,[15] of T4L. The ligand starts away from protein (unbound state) and unbiasedly samples the available conformational space, untill settled in binding pocket matching the crystal pose. (B) A typical example of conformational sampling of intrinsically disordered protein, shown here for sic1. In simulations, the proteins undergo/samples conformations as shown by representative snapshots.

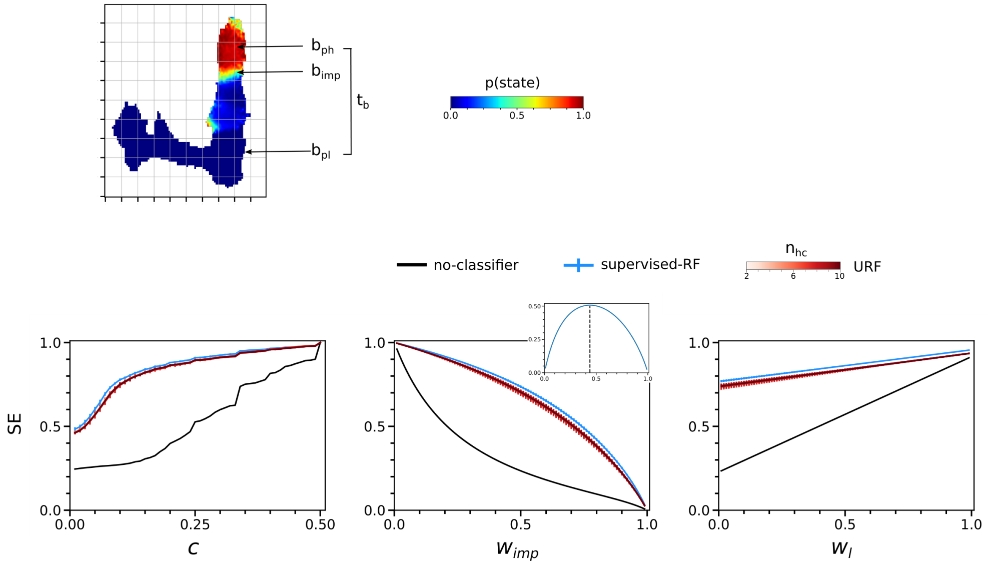


**Supplementary Figure 11:** Pictorial visualization of terms defined in methods (main). The behaviour of SE with respect to change in $c$, $w_{imp}$ and $w_l$. The inset in shows difference in supervised-RF and no-classifier, whose maxima represents the $w_{imp}$.

#### Vamp2 score

The vamp2 scores were estimated for each feature independently, as implemented in pyemma utility.[16,17] The input feature data was divided into train-test subsamples, with fitting on train data and reported vamp2 score was measured on test data. Since vamp2 score requires time series data, the random train-test subsampling of 0.7:0.3 ratio was not performed. For T4l constituting 6 trajectories, leave-one-out approach was utilized and repeated for six times. For mopR, cytochrome-P450 (channel-1) and Lennard Jones polymer constituting 129, 9 and 200 trajectories, 10 cross validation replicates were performed by randomly selected 100:29, 8:1 and 150:50 trajectories as train-test subsamples respectively. For cytochrome-P450 (binding) with only 3 trajectories, no train-test subsampling was performed. For sic1, phosphorylated-sic1, $\alpha$-synucien, ash1 and phosphorylated-ash1 constituting single trajectories, a random 0.2-0.3 fraction of end of trajectory was used as test subsample. The vamp2 scores was measures as varying lag times at a gap of 5 timesteps. The ones used for representation plots were performed at lag time same as tica lag time.

#### Train-loo

For 755 and 229 dimensional data of T4l and mopR, equal number of supervised random forest models were trained with leave-one-out (loo) scheme, i.e., trained on 754 and 228 dimensional input. Five fold cross-validation was performed by random 0.7:0.3 train:test splitting of data. The change in accuracy metric (T4l: f1 score, mopR: mean squared error) on test data compared to fully trained models (no leave-one-out) was used as feature importance score.

#### Predict-loo

For 755 and 229 dimensional data of T4l and mopR, equal number of prediction rounds were performed via leave-one-out scheme. For each round, a particular feature was set to (a) less-than-min or (b) more-than-max value. For instance, the 755 dimensional data of T4l with its range [0.23, 2.18], the less-than-min=0 and more-than-max=10 were used. The change in accuracy metric (T4l: f1 score, mopR: mean squared error) in two-round prediction was used as a measure of feature importance score. Five fold cross validations were performed by random 0.7:0.3 train:test splitting of data.

#### RFE

For recursive feature elimination (RFE), starting with all features, n (T4l: 37, mopR:21) rounds of elimination were performed with a number (T4L: 15, mopR: 9) of worst performing features (in supervised accuracy metric on test data) were discarded in each round, ultimately giving the best performing set of features (T4l: 200, mopR: 40). Five fold cross validations were performed by random 0.7:0.3 train:test splitting of data.

#### AMINO

Automatic mutual information noise omission (AMINO) was performed as defined in ref.[18] The max outputs (equivalent to number of important features or order parameters) was set to 200 and 40 for T4l and mopR respectively. The probability distribution was binned to 80 units for mutual information estimation. Five fold cross validations were performed by random 0.7:0.3 train:test splitting of data. The ultimate output yeilds selected set of order parameters decided according to distortion jumps (T4L: [199,199,192,198,192], mopR: [30,35,30,30,30]).

#### Spectral oasis

200 and 40 set of important features were selected via spectral oasis approach as defined in ref.[19] Spectral oasis approach performs a step-by-step selection of features with an aim to build a time-lagged covariance matrix similar to original data (full data). The *initial_columns* (number of features to start with) and *nsel* (new features to select at each step) were set to 1 and lag time was set to 700 (T4l) and 10 (mopR) steps (same as used in representation plots).

#### MoSAIC

The input 755 and 229 dimensional input of T4l and mopR ware clustered into groups of important features via correlation based feature selection strategy as implemented in MoSAIC (molecular systems automatic identification of cooperativity) package defined in ref.[20] The hyperparameters similarity metric, resolution parameter and weighted were set to correlation, 0.5 and True respectively. The ultimate output of MoSAIC was clusters of features arranged according to size (number of features) with the assumption that larger clusters corresponds to domains of important conformational changes. We have selected three cases: (a) features taken from the top cluster, (b) features taken from top 3 clusters and (c) features taken from all the clusters constituted by atleast 5 features. The three cases yields (a) T4l: [20,20,20,20,20], mopR: [205,206,206,205,205], (b) T4L: [49,49,49,49,49], mopR: [226,226,226,226,226], and (c) T4l: [127,128,128,127,128], mopR: [226,226,226,226,226,226] important features, in a five fold cross validation performed by random 0.7:0.3 train:test splitting of data.

#### SHAP

The shapely additive explanations (SHAP) approach was utilized to estimate feature importance scores as implemented in shap library and defined for MD systems in ref.[21] The supervised trained random forest model (on 0.7 train data) and test data (0.3 of input data) was used as input of shap algorithm which directly outputs the feature importance scores. As oppose to ref,[21] which utilizes committor values (estimated for predefined conformational states) as input labels, we directly use conformational states (or parameter defining it) as input labels. 200 and 40 important features were selected for T4l and mopR respectively.

#### Saliency-FNN/TVAE

Saliency scores, equivalent of feature importance scores in neural networks, were estimated by measuring the magnitude of the error gradient with respect to the input features, as defined in ref.[22] Two types of neural networks were employed: a feedforward neural network (FNN) and a time-lagged variational autoencoder (TVAE). The FNN was trained on supervised labels, where gradient is measured with respect to the mean squared error or binary cross-entropy between the predicted and target labels. In contrast, the TVAE was trained in an unsupervised manner to reconstruct input data, with the gradient measured against the reconstruction error. Multiple architecture and hyperparameter combinations were explored for both models (FNN: T4l=13, mopR=6; TVAE: T4l=5, mopR=6), and the final selections were made based on the reasonableness of their learning curvesÄîaccuracy for the FNN (ensuring no overfitting) and reconstruction error for the TVAE.

For the T4l dataset, the FNN was trained using 200 epochs, the Adam optimizer, binary cross-entropy loss, accuracy metrics, a batch size of 512, and ReLU activation functions (with sigmoid used for the final layer). Two architectures were finally used on standarized input data: one with a single node (755-1) and another deeper network (755-256-128-64-32-16-8-4-2-1) with an L1 regularizer (0.001) sequentially referred as architecture-1,2 in results. The TVAE for T4l were trained for 40 epochs using the Adam optimizer, a combination of mean squared error and KL divergence loss functions with a beta parameter of 0.001, a batch size of 32, and ReLU activations (with sigmoid in the final decoder layer). The input data were normalized such that the squared sum of each data instance equaled 1, and a lag time of 700 was used (same as used in representation plots). Several encoder-decoder architectures were explored sequentially referred as architecture-1-5 in results, i.e., 755-256-128-64-32-16-8-4-2, 755-256-64-16-4, 755-64-16-8-4, 755-64-32-16, and 755-32. The decoder architecture was same as encoder.

For the mopR dataset, the FNN was similarly trained for 200 epochs using the Adam optimizer, mean squared error as the loss function, a batch size of 128, and ReLU activations (sigmoid for the output layer). Six architectures were trained on standarized rescaled input data, 229-64-32-16-8-4-2-1, 229-32-8-1, 229-32-16-1, 229-16-16-1, 229-64-16-1, and 229-32-32-1. The mopR TVAE followed the same hyperparameters as the T4l TVAE, with 40 epochs, Adam optimizer, and the same loss function combination. The data preprocessing was also normalized to ensure a squared sum of 1 for each instance, with a lag time of 10 (same as used in representation plots). Six architectures were trained, i.e., 229-64-32-8-4-2, 229-32-8, 229-32-16, 229-16-16, 229-64-16, and 229-32-32. The decoder architecture was same as encoder.

Remaining hyperparameters were used as default in tensorflow library. Features were selected based on absolute magnitude of saliency scores (T4l:200, mopR:40). Five fold cross validations were performed by random 0.7:0.3 train:test splitting of data.

#### deepTICA

The input data of T4l and mopR was reduced to two best possible eigenvector components via deepTICA approach as defined in ref.[23] As opposed to single and biased trajectory used in ref.,[23] the data used in this work include multiple and unbiased trajectories. Hence, no time reweighing was performed and concatenated frame indices (different trajectories were separated by lag+3 gap) were used as time metric. The lag time of 700 (T4l) and 10 (mopR) steps were used, same as used for representation plots. The input data was rescaled to [0-1] and divided into 0.7:0.3 train:test subsamples. The architectures 755-256-64-16-2, 755-64-64-2, 755-32-32-4, 755-64-64-4 (architectures 1-4, T4l) and 229-128-64-32-8-2, 229-64-64-2, 229-32-32-2 (architectures 1-3, mopR) were trained with hyperparameters as follows: activ_type=tanh, loss_type=sum2 (maximizing eigenvalues of top 2 eigenvectors), num_epochs=200, lrate=0.001, l2_reg=0, and optimizer=adam. The training was stopped early (earlystop=True) if no improvement was achieved for 10 epochs (es_patience=10, es_consecutive=True, min_delta=0.001) and the best model was used.

#### AE-TICA

Autoencoders were used to dimensionally reduced the input data into lower dimensional latent space (T4l:200, mopR:40), equal to number of selected features via other approaches. Multiple architecture-hyperparameter (T4l:8, mopR:6) combinations were attempted and selected based on reasonability of learning curves (T4l:5, mopR:6). For T4l, architectures 755-215-256-200–755, 755-512-200–755, 755-400-200–755, and 755-200–755 (architectures 1-4) were trained with activation function relu. All architectures were also repeated for activation function tanh but only one could be trained reasonably (755-200–755, architecture-5). For mopR, architectures 229-128-64-40–229, 229-64-40–229, 229-40–229 were trained for activation functions relu (architectures 1-3) and tanh (arcitectures 4-6). The input data were scaled 0-1. The hyperparameters were set as follows: #epochs=200, batch_size=(T4l:512, mopR:128) loss=mean_squared_error, optimizer=adam, and kernel_initializer=glorot_uniform. The remaining hypearaprameters were used as default in tensorflow. The high dimensional latent space was further used in tica to generate representation plots.

#### TICA-VDE

TICA-VDE (time lagged independent component analysis - variational dynamics encoder) approach was used to dimensionaly reduced the input data (T4l, mopR) into two-dimensional latent space, as defined in ref.[24] The input data was first fed to TICA and resultant output was used as input to VDE model. VDE architectures with hidden_sizes=32 or 512 and hidden_depths=2 or 3 (architectures 1-4) were trained. The hyperparameters were set as follows: n_epochs=40, batch_size=1024, scale=0.001, dropout=0, learning_rate=0.0001, optimizer=adam, lag_time=(T4l:700, mopR:10 same as used for TICA), sliding_window=True, autocorr=True, loss=MSELoss (this also includes autocorrelation loss), activation=swish and remaining as default.

#### DiffNet

The functionally relevant conformational changes were identified by DiffNet as defined in ref.[25] DiffNet takes simulation ensembles in different states of a particular protein. For mopR, the apo-

WT, bound-WT, apo-MT and bound-MT (corresponding to data 3-6) simulation ensembles define its four states, and were used to train DiffNet. For T4l (data-1), the bound and unbound states were present within single simulation ensembles, hence cannot and were-not used to train DiffNet.

DiffNet is an autoencoder based architecture, where the encoder may be a single or double dense neural net which merged at the latent space. We used SAE type DiffNet (i.e., without splitting of encoder architecture), as the mopR's functionally important regions (allosteric linker) are far away from both binding and mutant (G148P) site. The $C\alpha$-aligned trajectories of mopR in its four states each constituting 37500 frames were used as input. DiffNet was trained on whitened 3d-coordinates of N, $C\alpha$, $C\beta$ and C atoms (except 148 residue to maintain same feature space across states). A single hidden layer of size 4x less than input feature space was used. In DiffNet, the latent layer was used in training for additional self-supervised classification task achieved via expectation-maximization algorithm. In this regard, the latent layer size was tested for 8, 32 and 64 to achieve best performance. The em_bounds were used as (0.1-0.4) and (0.6-0.9) for states corresponding to initial label 0 and 1 respectively. The initial labels were set as either [0,1,0,0] or [0,1,1,1] corresponding to apo-WT, bound-WT, apo-MT and bound-MT states. The training was performed till 10 epochs for each layer using batch size of 32 and learning rate of 0.0001. The rest of the hyperparameters were used as default. In addition to learning curves, the successful training was measured by three observables: (a) the reconstruction rmsd of aligned trajectories, (b) the DiffNet output labels, and (c) the morph output structures. For successfully trained DiffNet model, the aligned trajectories could be regenerated with low rmsd, the output labels corresponds to extremes (0 or 1) and morph structures were un-distorted.

For comparision with URF, the DiffNet was initially trained on whitened coordinates of all residues (1-229) but no successful training was observed for any hyperparameter set, potentially due to very high flexibility of terminal disordered regions. Hence, only residues 20-220 were considered. Finally, the successful training was performed for models with residues 20-220, act_map=[0,1,1,1] and latent size of 32 or 64, as reported in results.

#### Vampnet

For mopR (data-2) and $\alpha$-synuclien, the MSM equivalent metastable states were learned by vampnet as defined in ref.[26] Number of metastable states to be trained were kept similar to number of pcca metastable states in markov models, i.e., 3,4,5,6 for mopR and 2,3,4,5,6 for $\alpha$-synuclien. Vampnet consists of two neural network lobes which takes time lagged versions of same data as their input. Vampnets were trained at multiple lag times, similar to the ones used in MSM transition matrix lag times. Specifically, 10 and 14 different lag times were used for mopR (5,10,..50) and $\alpha$-synuclien (5,10,..70). Architecturally, both the network lobes were identical. The hidden layer architecture follows a continuous and constant decrease in number of neurons between successive layers by a constant factor, referred as *arc* in results (Fig. 4E). We used this factor as 2,3,4 and 6 to encompass different possible hyperparameter space. For instance, for a mopR vampnet model having 3 metastable states, 229 input dimension and a factor of 2, the hidden layer architecture shall be 229-96-48-24-12-6-3. Similarly, other architectures were designed in accordance with original work.

For each neural network lobe, random dropout of 0.1 was used in initial layers. For deeper network, first two layers were subjected to dropout, while only first layer for shallower networks. Specifically for one very shallow architecture of mopR (229-36-6, number of metastable states=6, factor=6) no dropout was performed. Learning rate and epsilon were used as $10^{-3}$ and $10^{-}5$ respectively. Activation function of relu for all layers and softmax for last layer were used. Initially the pre-training was performed for 50 epochs with vamp1 score as loss function. Final training for 100 epochs was performed with vamp2 score as loss function. The batch size of 4096 was used.

The input data was scaled as per standard deviation around the mean. Five-fold cross validation was performed with 0.7:0.3 random train:test subsampling. Overall, by varying lag time, number of metastable states, network architecture and cross validation, 800+1400 vampnet models were trained for mopR and $\alpha$-synuclien respectively. The successful training was confirmed by learning curves (Fig S) and vampnet models with unstable learning curves were discarded. Further, vampnet models with zero population of one or more metastable states (i.e., unable to learn the desired number of metastable states) were also discarded. Overall most models were successfully trained, only 4 vampnet models were discarded for mopR and none for $\alpha$-synuclien.

**Supplementary Table 1:** Python libraries used in this work. The versions may be different for baseline approaches, as per specific requirements.

| | |
|---|---|
| Python | 3.9.12 |
| numpy | 1.25.1 |
| matplotlib | 3.7.2 |
| pyemma | 2.5.12 |
| sklearn | 1.3.0 |
| shap | 0.45.0 |
| tqdm | 4.63.0 |
| scipy | 1.11.1 |
| sys | 3.9.12 |
| keras | 2.15.0 |
| tensorflow | 2.15.0 |
| MDAnalysis | 2.5.0 |
| numba | 0.59.0 |
| fastcluster | 1.2.6 |

(5) Roy, R.; Ray, S.; Chowdhury, A.; Anand, R. Tunable multiplexed whole-cell biosensors as

environmental diagnostics for ppb-level detection of aromatic pollutants. *ACS sensors* **2021**, *6*, 1933–1939.
